# Astrocyte CB_1_ receptors drive blood-brain barrier disruption in CNS inflammatory disease

**DOI:** 10.1101/2025.06.23.660998

**Authors:** T Colomer, E Sánchez-Martín, A Bernal-Chico, A Moreno-García, R Serrat, AM Baraibar, A Uribe-Irusta, A Iriarte-Sarria, U Skupio, C Matute, V Tepavcevic, I Fernández-Moncada, C Chapouly, G Marsicano, S Mato

## Abstract

Reactive astrocytes shape central nervous system (CNS) inflammation and participate in myelin damage and repair mechanisms in multiple sclerosis (MS). Through the activation of cannabinoid CB_1_ receptors (CB_1_R) expressed by neurons and oligodendrocyte lineage cells, endocannabinoid signaling restricts neurodegeneration and promote remyelination in preclinical MS models. However, despite accumulating evidence that supports a crucial role for these receptor populations in brain physiology and pathology, the implications of astrocyte CB_1_R signaling in MS initiation and progression remain uncertain. Using complementary *in vivo* disease models, here we investigated the effects of targeted genetic deletion of astrocytes CB_1_R on the expression of MS-like pathology in mice. Interestingly, astrocyte-specific deletion of CB_1_R reduced demyelinating neuropathology, attenuated astrocyte reactivity and improved clinical deficits during the time-course of experimental autoimmune encephalomyelitis (EAE). Mice with astrocyte CB_1_R inactivation displayed unaltered oligodendrocyte populations both in EAE lesions and in lysolecithin-induced remyelinating spinal cord lesions, likely excluding that astrocyte CB_1_R modulate myelin repair processes. Conversely, inactivation of CB_1_R in astroglial cells restricted humoral and leukocyte parenchymal infiltration and reduced the expression of vascular effectors in EAE lesions. Finally, loss of blood-brain barrier (BBB) function induced by cortical microinjection of VEGF-A was less severe in GFAP-CB_1_R-KO mice. These results show that astrocyte CB_1_R signaling constitutes a significant pro-inflammatory mechanism in MS and bring to light a deleterious role for endocannabinoid-mediated modulation of astroglial cells with potential implications in the etiopathology and therapy of neuroinflammatory disorders.

## Introduction

Multiple sclerosis (MS) is an immune-mediated inflammatory disease of the central nervous system (CNS) and one of the most prevalent neurological disorders leading to chronic disability among young adults (1). The main pathological hallmark of MS is the formation of demyelinating lesions in the brain and spinal cord associated to neuroaxonal degeneration as primary substrate of the irreversible clinical deficits that characterize disease progression (2). Focal lesions in MS are thought to be caused by the bidirectional interaction between peripheral immune cells that infiltrate into the CNS parenchyma, including T cells, B cells and myeloid cells, and activated resident immune cells, mainly astrocytes and microglia (3). A subset of lesions in MS patients are characterized by a variable extent of remyelination suggested as a mechanism of neuroprotection and clinical recovery (4, 5). At present, treatment strategies for clinical exacerbation in MS include almost exclusively immunotherapeutic drugs that target peripheral immune cells and their trafficking into the CNS and lead to a substantial reduction in lesion formation and clinical relapse rate but do not prevent the progression of clinical disability.

Astrocytes support brain homeostasis and function through a plethora of mechanisms that include the modulation of synaptic transmission through the release of gliotransmitters, the metabolic assistance to neurons and oligodendrocytes and the maintenance of the blood-brain barrier (BBB), among others. In recent years, complementary lines of evidence have shown that astrocytes adopt a wide spectrum of reactive states during neuroinflammation that confer these cells the potential to exacerbate damage or facilitate repair (6–8). Aberrant astrocyte activation critically contributes to inflammatory lesion formation and progression in MS through the release of molecules that promote the loss of BBB integrity and the recruitment of peripheral immune cells (8–10). However, astrocytes may also facilitate the differentiation of oligodendrocyte progenitor cell (OPCs) and support the survival of mature oligodendrocytes to ensure successful remyelination (11, 12). An improved understanding of astrocyte regulatory mechanisms in neuroinflammatory contexts may thus provide novel therapeutic targets that reduce MS pathology and clinical severity during acute and progressive phases of the disease.

Cannabinoid type-1 receptor (CB_1_R) is one of the most abundant G protein-coupled receptors in the mammalian brain and the primary molecular target of endogenous cannabinoids - anandamide and 2-arachidonoylglycerol - and Δ^9^-tetrahydrocannabinol (THC), the main psychoactive component of the hemp plant *Cannabis sativa* (13). Endocannabinoids acting on CB_1_R heterogeneously expressed in neuronal populations and glial cells modulate physiological brain functions through a wide variety of mechanisms and exhibit neuroprotective potential during CNS damage (14–16). Pharmacological and genetic studies conducted so far have demonstrated that cannabinoid agents that potentiate CB_1_R-mediated signaling attenuate neurodegeneration and suppress neuroinflammation while promoting myelin repair in rodent models of MS (17–20). However, the clinical efficacy of cannabinoid-based medications in MS is limited, and therapeutic optimization requires a better understanding of endocannabinoid and CB_1_R related networks in inflammatory demyelinating contexts (16, 21). On mechanistic grounds, there is consensus that the neuronal population of CB_1_R provides neuroprotection from excitotoxic damage in rodent models of MS by limiting synaptic glutamate release (20, 22–24). Experimental *in vivo* evidence from conditional mutant mice has also recently grounded the hypothesis that CB_1_R expressed by OPCs promote oligodendrocyte differentiation and favor remyelination in MS (25, 26). However, the implications of CB_1_R in astrocytes during MS onset and progression have been largely neglected despite the fundamental contribution of these receptor pools as effectors of (endo)cannabinoid-mediated signaling in the brain (27, 28). Indeed, CB_1_R activity has arisen during the last decade as crucial modulator of astrocyte-derived gliotransmitter release and metabolic supply with emerging implications in neuroinflammatory disorders (29, 30)

In this study, we aimed at disentangling the roles of astrocyte CB_1_R (aCB_1_R) signaling in MS neuropathology using mice where this receptor population is selectively ablated in astroglial cells and a combination of rodent models that recapitulate autoimmune demyelination, BBB breakdown and myelin repair. Our results highlight that aCB_1_R exacerbates neurological disability during autoimmune demyelination by fostering BBB permeability and recruitment of peripheral immune cells towards lesion sites. These observations identify a previously unexpected pathogenic role of aCB_1_R during CNS inflammatory lesion formation with potential implications in MS pathogeny and therapy.

## Materials and methods

### Mice

All experiments were performed in accordance with the Guide for the Care and Use of Laboratory Animals (National Research Council Committee, 2011) and the European Communities Council Directive of 22 September 2010 (2010/63/EU74). Experiments were approved by the local ethical committees of the University of the Basque Country (approval numbers 2017140, 2020005 and 2022245) and the University of Bordeaux (approval numbers A33063098 and 16901). Inducible mutant mice lacking CB_1_R in cells expressing the astrocytic maker glial fibrillary acid protein GFAP (aCB_1_-KO) and aCB_1_-WT littermates bred at the Neurocentre Magendie (Bordeaux, France) were used. Cages were enriched and mice were maintained under standard conditions (food and water *ad libitum*; 12 h-12 h light-dark cycle). Experiments were performed during dark cycle (light off at 8:00 h a.m.). *In vivo* models were induced in female mice based on epidemiological evidence that MS affects 2-4 times more women than men (31). The number of mice in each experimental group was similar. No statistical methods were used to predetermine sample size.

aCB_1_-KO mice were generated by crossing CB_1_^f/f^ mice (32) with GFAP-CreERT2 mice (33) using a three-step backcrossing procedure to obtain CB_1_^f/f;GFAP-CreERT2^ and CB_1_^f/f^ littermates, called aCB_1_-KO and aCB_1_-WT respectively. Deletion of the *Cnr1* gene was obtained in adult mice (6-12 weeks of age) by daily intraperitoneal (i.p.) injections of tamoxifen (1 mg dissolved at 10 mg/mL in 90% sesame oil, 10% ethanol) for 8 days (34, 35). Mutant mice were used for experiments 3-4 weeks after the last tamoxifen injection.

### EAE model

Mice were immunized in the flank by subcutaneous (s.c.) injection of 200 μg MOG_30-55_ peptide (MEVGWYRSPFSRVVHLYRNGK) (Peptide Synthesis Core Facilities of the Pompeu Fabra University, Spain) in incomplete Freund’s adjuvant supplemented with 8 mg/mL Mycobacterium tuberculosis H37Ra (Difco Laboratories). Pertussis toxin (500 ng; Calbiochem) was injected i.p. on the day of immunization and again 2 days later. Body weight and motor symptoms were recorded daily and scored from 0 to 8 as follows: 0, no detectable changes in muscle tone and motor behavior; 1, flaccid tail; 2, paralyzed tail; 3, impairment or loss of muscle tone in hindlimbs; 4, hindlimb hemiparalysis; 5, complete hindlimb paralysis; 6, complete hindlimb paralysis and loss of muscle tone in forelimbs; 7, tetraplegia; and 8, moribund.

### Demyelinating lesion induction

Demyelinating lesions were induced stereotaxic injection of 0.5 μL of 1% lysophosphatidylcholine (LPC) (Sigma-Aldrich) diluted in sterile saline solution (0.9% NaCl) in the spinal cord (36). Mice were anesthetized by i.p. injection of a ketamine (100 mg/Kg; Fatro) and xylazine (20 mg/Kg; Calier) cocktail. Two longitudinal incisions into the *longissimus dorsi* at each side of the vertebral column were performed and the muscle tissue covering the column was removed. Animals were placed in a stereotaxic frame and the intervertebral space of the 13th thoracic vertebra was exposed by removing the connective tissue. An incision into dura mater was performed using a 30-gauge needle and LPC was injected into the white matter of the *dorsal funiculus* via a Hamilton syringe attached to a glass micropipette using a stereotaxic micromanipulator. The lesion site was marked with sterile charcoal. Following LPC injection, the muscle sheaths were closed using 3/0 Monocryl and the wound was sutured with 4/0 silk. Buprenorphine (0.1 mg/Kg; Dechra) was subcutaneously administered as postoperative analgesic treatment. Mice were euthanized and processed for immunohistochemistry 14 days after the surgery.

### VEGF-A microinjection

Mice were anesthetized using a local injection of lidocaine (20 mg/mL) under the skull skin and a mix of air and isoflurane (3% for induction and 1.5% for support). Animals were placed into a stereotactic frame and mouse VEGF-A_165_ (60 ng in 3 μL NaCl 0.9%) or vehicle were delivered into the cerebral cortex at y = 1 mm caudal to Bregma, x = 2 mm, z = 1.5 mm as previously described (37). Mice received a subcutaneous injection of buprenorphine (0.1 mg/Kg) (Vetergesic) 30 minutes before surgery and again 8 hours post-surgery to assure constant analgesia. Animals were sacrificed 24-48 h after VEGF-A_165_ injection and processed for immunohistochemistry.

### Surgery for AAV administration and fiber implantation

Mice were anaesthetized with isoflurane and placed on a heating-pad to keep the body temperature at 37°C. Eye dehydration was prevented by topical application of ophthalmic gel and analgesia was achieved by s.c. injection of buprenorphine (Buprecare, 0.05 mg/Kg). The skin above the skull was shaved with a razor and disinfected with modified ethanol 70% and betadine before an incision was made. Mice were placed into a stereotaxic apparatus (David Kopf Instruments) with mouse adaptor and lateral ear bars and injected with an AAV encoding the genetically encoded Ca^2+^ indicator GCamp6s under the GFAP promoter (AAV-9/2-GFAP-hHBbI/E-GCaMP6s-bGHp(A)), to carry out fiber photometry imaging of Ca^2+^ activity in astrocytes. Virus titers were between 10^10^-10^12^ genomic copies per mL. Stereotaxic injections were targeted to the mouse somatosensory cortex according to the following coordinates (from bregma): anterior-posterior −1.5; medial-lateral ± 2.5; dorsal-ventral −1.5. Viral particles were injected at 400-500 nL alone or in combination at a maximum rate of 100 nL/min using a glass pipette attached to a Nanojet III (Drummond, Broomall, USA). Following virus delivery, the syringe was left in place for 10 min before being slowly withdrawn from the brain. The optical fiber (400 μm diameter) was placed 250 µm above the injection site during the same surgical session. Mice were weighed daily and individuals that failed to return to their pre-surgery body weight were excluded from subsequent experiments. Mice were treated with tamoxifen 1 week after the surgery.

### Fiber photometry imaging

Freely moving mice were imaged after 3 days of handling habituation. The day of recording each mouse was placed in a rectangular chamber and its behavior recorded using a camera placed above the chamber. Baseline recordings of spontaneous astrocyte activity were made for 15 min every 1-2 days starting 3 days before MOG administration. The calcium signal evoked by sensory stimulation of the tail was assessed at the end of the baseline period.

Cortical astrocyte GCaMP6s were imaged *in vivo* using 470 and 405 nm LEDs. The emitted fluorescence is proportional to the calcium concentration for stimulation at 470 nm (38, 39). The isosbestic 405 nm stimulation (UV light) was used in alternation with the blue light (470 nm) for analysis purposes as the fluorescence emitted after this stimulation is not depending on calcium (40). The GCaMP6s fluorescence from the astrocytes was collected with a sCMOS camera through an optic fiber divided in 2 sections: a short fiber implanted in the brain of the mouse and a long fiber (modified patchcord), both connected through a ferrule-ferrule (1.25 mm) connection. MATLAB program (Matlabworks) was used to synchronize each image recording made by the camera, and the GCaMP6s light excitation made by the LEDs (470 and 405 nm). The two wavelengths of 470 and 405 nm at a power of 0.1 mW were alternated at a frequency of 20 Hz each (40 Hz alternated light stimulations).

To calculate fluorescence due specifically to calcium fluctuations and to remove bleaching and movement artifacts, the isosbestic 405 nm signal was subtracted from the 470 nm calcium signal. Specifically, normalized fluorescence changes (Δ*F*/*F_0_*) were calculated by subtracting the mean fluorescence (2 min sliding window average) from the fluorescence recorded by the fiber at each time point and dividing this value by the mean fluorescence (*F-F*_mean_)/*F*_mean_) using a customized Matlab software. Subsequently, the calcium independent isosbestic signal was subtracted to the raw signal emitted after the 470 nm excitation to eliminate unspecific fluorescence. The result will be the global calcium signal (Δ*F*/*F* (%) = Δ*F*_Ca_ - Δ*F*_iso_), that was used as an estimate of tonic activity of the astrocytes. Calcium transients were detected on the filtered trace (high filter) using a threshold to identify them (2 median absolute deviation -MAD-of the entire trace). Duration and frequency were calculated on the detected transients. Amplitude was determined as the MAD of each studied period (41).

### Quantitative RT-PCR

Mice were anesthetized with ketamine/xylazine (80/10 mg/Kg, i.p; Imalgene^®^/Rompun^®^) and transcardially perfused with cold phosphate buffer saline (PBS) for 30 s in order to remove blood cells from the brain. The lumbar spinal cord was dissected in lysis buffer containing 1% β-mercaptoethanol for optimal template preservation. Total RNA was purified with on-column DNAse treatment using RNeasy Plus Mini kit (Qiagen; 74104) following manufacturer’s instructions. RNA was eluted with 14-35 μL of RNAse-free deionized water and stored at −80°C until analysis. Synthesis of cDNA, pre-amplification and amplification steps were performed at the Genome Analysis Platform of the UPV/EHU following quality control of RNA samples with an Agilent 2100 Bioanalyzer (Agilent Technologies). Pre-amplified cDNA samples were measured with no reverse transcriptase and no template controls in the BioMark HD Real-Time PCR System using 48.48 Dynamic Arrays of integrated fluidic circuits (Fluidigm Corporation). We used commercial primers from IDT Integrated DNA Technologies or Fluidigm Corporation (**Supplementary Table 1**). Data pre-processing and analysis were completed using Fluidigm Melting Curve Analysis Software 1.1.0 and Real-time PCR Analysis Software 2.1.1 (Fluidigm Corporation) to determine valid PCR reactions. *Gapdh*, *Hprt* and *Ppia* were included as candidate reference genes for normalization purposes. Data were corrected for differences in input RNA using the geometric mean of reference genes selected according to results from the normalization algorithms geNorm (https://genorm.cmgg.be/) and Normfinder (https://moma.dk/normfinder-software). Relative expression values were calculated with the 2^-ΔΔCt^ method.

### Western blot

Anesthetized mice were transcardially perfused with cold PBS and the lumbar spinal cords were homogenized (1:20 w/v) in ice-cold RIPA buffer (Thermo Scientific™; 89900) containing a protease inhibitor cocktail (Thermo Scientific™; 87786) by using a Potter homogenizer provided with a loosely fitting Teflon pestle. Samples were incubated in ice for 30 min and subjected to centrifugation (12000 *x* g at 4°C for 8 min) to remove insoluble material. Solubilized proteins were quantified in the supernatants using the BioRad Protein Assay Kit (Protein Assay Reagents; 5000-114-13-15). Protein samples (4 µg) were loaded into polyacrylamide Criterion TGX Precast (BioRad) gels before electrophoretic transfer onto Nitrocellulose membranes (Amersham™ Protran^®^ Western blotting membranes, pore size 0.2 μm). Membranes were blocked for 1 h in Tris-buffered saline (TBS) (50 mM Tris, 200 mM NaCl, pH 7.4) with 0.05% Tween-20, 5% BSA. Subsequently, membranes were incubated overnight at 4°C with primary antibodies raised against myelin basic protein (MBP; 1:1000; BioLegend; 808401), Claudin 4 (CLN-4; 1:500; ThermoFisher; 32-9400), Cadherin 5 (CDH-5; 1:500; R&D Systems; AF1002), Intercellular Adhesion Molecule 1 (ICAM-1; 1:500; R&D Systems; AF796), Podocalyxin (PODXL; 1:500; R&D Systems; AF1556), Vascular cell adhesion protein 1 (VCAM-1; 1:1000; Abcam; ab134047), zona occludens (ZO-1; 1:1000; Invitrogen; 02200) and α-Tubulin (1:5000; Abcam; ab7291) in blocking solution. Membranes were incubated for 1 h at RT with horseradish peroxidase-conjugated secondary antibodies (1:5000; Cell Signaling Technology) and developed with NZY Standard ECL Western Blotting Substrate (NZYtech). Volumetric analysis of relevant immunoreactive bands was carried out after acquisition on a ChemiDoc XRS System (Bio-Rad) controlled by The Quantity One software v 4.6.3 (BioRad).

### Histology and fluorescence immunohistochemistry

#### Mice sacrifice and tissue processing

EAE mice were deeply anesthetized by intraperitoneal injection of Dolethal (200 mg/Kg) and perfused with 4% paraformaldehyde (PFA) in 0.1 M phosphate buffer (PB) (25 mM NaH_2_PO_4_·H_2_O, 75 mM Na_2_HPO_4_; pH 7.4) for 10 min. The spinal cords were extracted and post-fixed for 24 h at 4°C in the same fixative solution. Alternatively, anesthetized mice were transcardially perfused with cold PBS and the brains extracted and stored at −20°C. Mice injected with LPC were perfused with 2% PFA in 0.1 M PB for 15 min and spinal cords post-fixed for 30 min. Spinal cord tissues from EAE and LPC-injected mice were cut into 1-4 longitudinal 2-mm-thick blocks and placed in 15% sucrose-7% gelatin in PBS. Tissue blocks were then frozen in isopentane for 2 min at −65°C and stored at −80°C. Coronal 10-12 μm-thick spinal cord and forebrain sections were cut into Superfrost glass slides (Thermo Fisher, #11976299) using a CM3050 S cryostat (Leica Biosystems) and stored at −20°C. Demyelination and glial reactions were evaluated using Luxol Fast Blue (LFB) and immunofluorescence standard protocols.

Mice injected with VEGF-A received a subcutaneous injection of buprenorphine (0.1 mg/kg) (Vetergesic) 30 min before surgery. Mice were then profoundly anesthetized by i.p. injection of a mix of ketamine (100 mg/kg) (Imalgene) and xylasine (20 mg/kg) (Rompun), and transcardially perfused with PBS for 5 min and then with 10% formalin (Merck, #252549) for 12 min. The brains were post-fixed in 10% formalin for 3 h and incubated in 30% sucrose overnight. Forebrain tissues were embedded on Tissue-Tek O.C.T.Compound (Sakura, #4583) and stored at −80°C. Coronal 12 μm-thick sections containing the marked lesion were cryostat sectioned into Superfrost glass slides and stored at −20°C until further processing.

#### Immunolabelling of spinal cord tissue

Spinal cord sections were air-dried for 30 min at RT and rehydrated in Tris buffer saline (TBS) (20 mM Tris, 1.4 M NaCl; pH 7.6) for 30 min. Antigen retrieval was performed for Olig2 immunostaining by adding low-pH R-Universal retrieval buffer (Aptum Biologics, #AP0530-500) and heating the slices in a microwave for 45 s at maximum temperature. For MBP immunolabelling, slices were permeabilized in absolute ethanol for 15 min at −20°C followed by extensive washing in TBS. Tissue sections were incubated for 1 h at RT in blocking solution containing 5-10% normal goat serum (NGS) (Vector Labs, S-1000) or donkey serum (NDS) (Interchim, #UP77719A K) and 0.2% Triton X-100 in TBS. The blocking solution was supplemented with 3% Fab fragment (Jackson ImmunoResearch) when using primary antibodies made in mouse. Slides were incubated overnight at 4°C with primary antibodies (**Supplementary Table 2**) diluted in TBS containing 5% NGS and 0.1% Triton X-100, washed in TBS (3 × 10 min) and incubated for 1 h at RT with Alexa Fluor secondary antibodies made in goat or donkey and Hoechst (4 μg/mL) (Sigma-Aldrich; B2261) in antibody solution. Tissue sections were washed with TBS (3 × 10 min) and mounted with ProLong Gold Antifade (Thermo Fisher; P36930), or Fluoromount-G (Thermofisher, 00-4959-52) mounting media for microscopy analysis.

#### Luxol fast blue myelin staining

Spinal cord sections were incubated overnight at 37-42°C with 0.1% Luxol fast blue (LFB) (Sigma-Aldrich, 1328-51-4) diluted in 95% ethanol and 0.5% glacial acetic acid and subsequently differentiated with a 0.01% lithium carbonate solution. Tissues were dehydrated by immersion in increasing concentrations of ethanol and the processed with xylene and mounted with DPX mounting medium.

#### Immunolabelling of cortical tissue

Cryostat sections from EAE mice containing the somatosensory cortex were air-dried for 30 min at RT, fixed in 4 % PFA for 15 min and washed in TBS (3 × 10 min). Tissue slides were incubated for 1 h at RT in a blocking-permeabilization solution containing 5% NGS and 0.2% Triton X-100 in TBS and subsequently incubated for 12-48 h at 4°C with primary antibodies (**Supplementary Table 2**) diluted in blocking solution. Following extensive washing, primary antibodies were detected by incubation with appropriate Alexa Fluor secondary antibodies for 2 h at RT. Hoechst was used for chromatin staining. Sections were then washed in TBS (3 × 10 min) and mounted in ProLong mounting medium using coverslips for microscopy analysis.

Tissue sections from VEGF-A-injected mice were tempered for 20 min and then rehydrated in PBS for 20 min. Antigen retrieval was performed by incubation in a Tris-EDTA solution (10 mM Tris HCl, 1 mM EDTA; pH 8) for 30 min at 100°C. Slices were then washed in PBS (3 x 5 min) and incubated for 1 h at RT in a blocking solution containing 10% NDS and 0.3% Triton X-100 in PBS. Sections were subsequently incubated overnight at 4°C with the corresponding primary antibodies (**Supplementary Table 2**) prepared in 5% NGS and 0.1% Triton X-100 in PBS. Tissue sections were washed in PBS (3 x 5 min) and incubated with Alexa Fluor secondary antibodies for 1 h at RT. Following extensive washing tissues were mounted on glass coverslips using Fluoromount-G with DAPI mounting medium (Thermofisher, # 00-4959-52) for microscopy analysis.

#### Image acquisition and analysis

Quantitative analysis of demyelination and inflammatory lesion number in EAE mice was performed in 2-4 tissue sections per mouse imaged using a 3D Histech Panoramic MIDI II slide scanner and the CaseViewer and CaseConverter softwares (3DHistech). For the immunohistochemical characterization of brain and spinal cord tissues, optical images from sections processed in parallel were acquired using 20X and 40X lens on a Leica TCS STED CW SP8 super-resolution microscope or a Zeiss Axioplan 2 pseudoconfocal microscope coupled to an Axiocam MRc5 digital camera. Image acquisition was carried out using fluorescence intensity settings at which the control sections without primary antibody gave no signal.

Immunohistochemical characterization of demyelinating lesions in the EAE model was performed by examining 4 objective pictures taken from 2 non-consecutive spinal cord sections per mouse. Regions of interest (ROIs) corresponding to the demyelinated plaque and the adjacent periplaque were established through MBP immunohistochemistry and the density of the cell nuclei. The plaque of demyelinated lesions was characterized by the total lack of MBP immunostaining and a high nuclear density, while the periplaque was determined as the area corresponding to a 100 μm perimeter measured from the lesion edge to the adjacent area, and characterized by weak or less dense MBP immunostaining. Immunolabelling of LPC injected spinal cords was evaluated in 2 tissue sections containing the central part of the demyelinating lesion. The somatosensory cortex of EAE mice was examined for inflammatory lesion load and astrocyte reactivity and immunohistochemical characterization performed in 2-4 images collected from 2 non-consecutive coronal sections per mouse. Cortical lesions induced by VEGF-A were evaluated in 3-4 images collected from the lesion area.

Image analysis was performed using Fiji Image J (42). Immunopositive cells were counted in a selected region of interest (ROI) using a cell counter plugin and data expressed as mean cell number per square millimeter (mm^2^) of tissue area. Analysis of immunostained areas was performed in 16-bit gray scale transformed pictures. Fluorescence signals were considered positive if they were above a defined intensity threshold and normalized to total selected ROI area. For colocalization analysis, pixels positive for GFAP or Iba-1 and C3, ICAM-1, VCAM-1, VEGF-A or CD68 immunoreactivity were counted in projections of Z series stacks with the same number of images taken at a spacing of 0.8 μm by a blinded observer.

### Data collection and statistical analyses

No statistical methods were used to pre-determine sample sizes but they are similar to those reported in previous publications. Experimenters were always blinded to mice genotype but not to treatments. Statistical analyses were performed using GraphPad Prism 10 for Windows (GraphPad Software Inc). Summary results are presented as the mean of independent data points ± SEM. Datasets were initially tested for normal distribution with the Shapiro-Wilk test and differences between groups were determined by two-tailed unpaired Student *t* test, Mann-Whitney test or two-way ANOVA followed by Šídák’s test for multiple comparisons. Differences were considered to be significant when *p* < 0.05.

## Results

### Astrocyte CB_1_ receptors exacerbate clinical deficits and myelin pathology in EAE

To study the role of aCB_1_R in MS we analyzed the phenotype of conditional mutant mice lacking CB_1_R in GFAP positive cells (Han et al., 2012) in the EAE model of autoimmune demyelination. Upon EAE induction, mice lacking CB_1_R specifically in astrocytes (aCB_1_-KO) displayed similar disease onset but significantly decreased clinical scores during the acute phase of the disease (**Figure 1a**). Histological evaluation of spinal cords at the experimental end-point (22 dpi) revealed that the number of inflammatory lesions, mostly associated to white matter areas close to the tissue edge, was reduced in aCB_1_-KO mice as compared to aCB_1_-WT controls (**Figure 1b**). Consistently, immunostaining for the myelin protein MBP showed attenuated demyelination in spinal cord tissue sections from aCB_1_-KO mice (**Figure 1c**).

**Figure 1.**
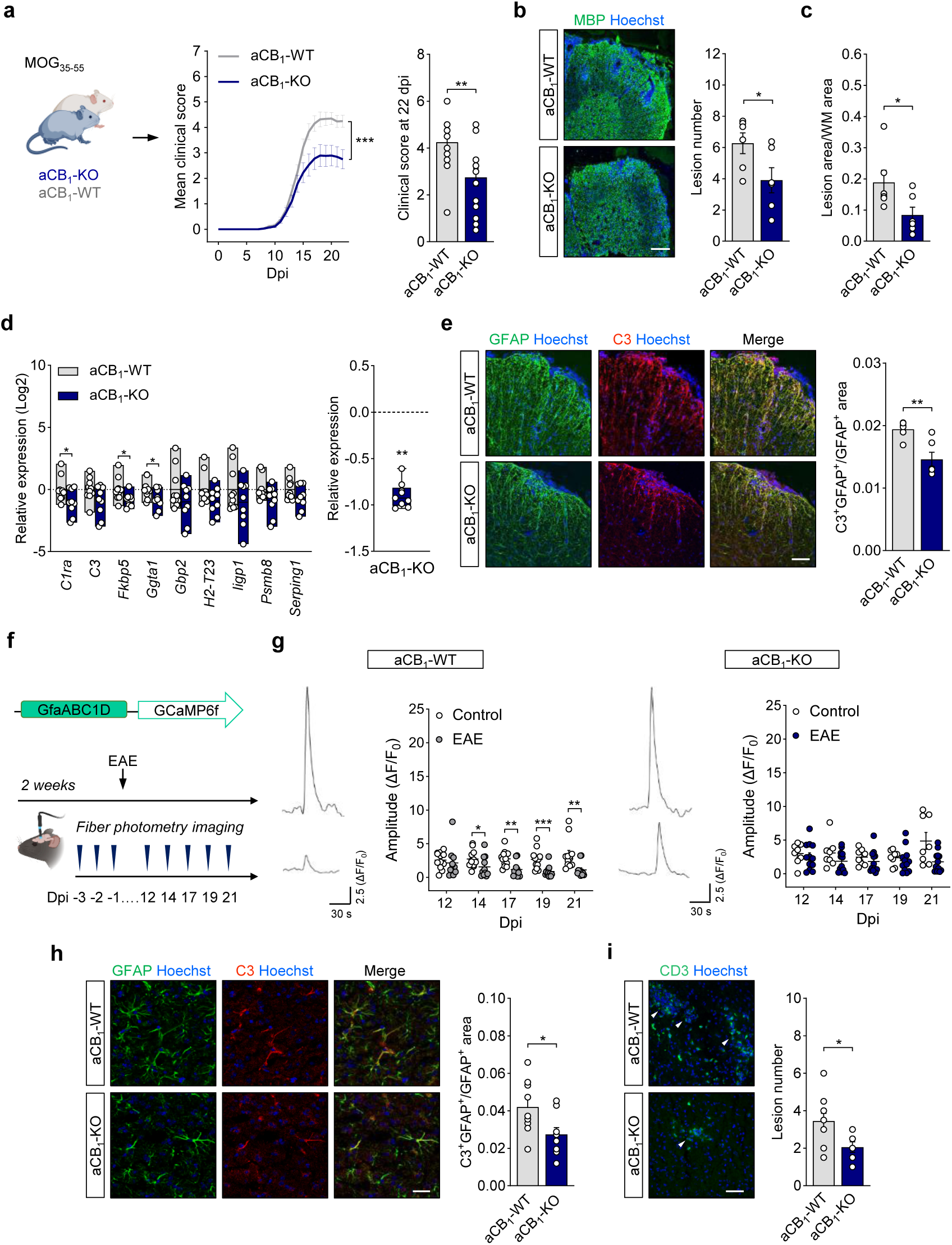
Astrocyte CB_1_R exacerbates EAE pathology and clinical deficits. (**a**) Time-course of EAE progression in aCB_1_-KO mice and aCB_1_-WT littermates. Comparison of motor scores revealed significantly attenuated neurological disability during acute EAE disease. Data are representative of 2 independent EAE experiments pooled together (*n* = 18-19 mice). (**b**-**c**) Representative images of MBP immunostaining in spinal cord sections from aCB_1_-KO and aCB_1_-WT mice at 25 days post-immunization (dpi). Scale bar = 200 µm. Quantitative analysis shows reduced lesion load (**b**) and demyelinated area (**c**) in aCB_1_-KO mice (*n* = 6 mice). (**d**) Relative expression of neurotoxic astrocyte genes in spinal cord tissue from aCB_1_-KO and aCB_1_-WT mice at 22 dpi (*n* = 9-10 mice). *Right*: Comparative analysis of neurotoxic astrocyte gene expression profile between aCB_1_-KO and aCB_1_-WT mice (*n* = 9 genes). (**e**) Representative confocal z-stack projections depict immunolabeling of the astrocyte reactivity marker GFAP and complement component 3 (C3) in spinal cord tissue from aCB_1_-KO and aCB_1_-WT mice. Scale bar = 100 µm. Quantitative analysis of GFAP and C3 colocalization shows attenuated C3 in astroglial profiles from aCB_1_-KO animals (*n* = 6 mice). (**f**-**i**) Attenuated cortical astrocyte pathology and functional deficits in aCB_1_R null mice during EAE. (**f**) Experimental approach for *in vivo* time-course analysis of astrocytic calcium activity during EAE. Fiber photometry imaging was performed in aCB_1_-KO and aCB_1_-WT mice recorded in parallel for 3 consecutive days before EAE induction by immunization with MOG_30-55_ and at 12, 14, 17, 19 and 21 dpi. (**g**) aCB_1_-KO mice show preserved astrocyte network function during acute EAE disease. Representative traces show calcium responses of cortical astrocytes evoked by tail-holding in aCB_1_-WT (*left*) and aCB_1_-KO (*right*) mice at −3 (*top*) and 19 (*bottom*) dpi. Graphs depict the amplitude of cortical astrocyte calcium responses recorded from aCB_1_-WT (*left*) and aCB_1_-KO (*right*) during the EAE time course compared to values from non-immunized animals recorded in parallel (*n* = 8-12 mice). (**h**) Representative confocal micrographs and quantification of C3 immunoreactivity in GFAP immunopositive astrocyte profiles within somatosensory cortex layer V-VI from aCB_1_-KO and aCB_1_-WT mice at acute EAE disease (22 dpi; *n* = 8-9 mice). Scale bar = 25 µm. (**i**) Confocal micrographs and quantification of CD3 immunopositive T inflammatory lesions (arrowheads) in cortical tissue from aCB_1_-KO and aCB_1_-WT mice (*n* = 8-9 mice). Scale bar = 50 µm. **a**, \*\*\**p* < 0.0001, Wilcoxon signed rank test for the comparison of score curves from the onset of EAE symptoms at 8 dpi to 22 dpi and \*\**p* = 0.0073, Mann-Whitney test. **b**-**i**, \**p* < 0.05, \*\**p* < 0.01 and \*\*\**p* < 0.001, unpaired *t*-test or Mann-Whitney test.

We next interrogated the protective phenotype of aCB_1_R-KO mice at chronic EAE disease stages. Astrocyte-specific CB_1_R null mice displayed a sustained reduction in disability scores during EAE progression to more chronic clinical plateau (**Figure S1a**). Comparative analysis at 35 dpi evidenced a non-significant attenuation of motor symptomatology in aCB_1_-KO mice that was associated to improved spinal cord myelin pathology in terms of demyelinating lesion numbers and proportion of demyelinated white matter area (**Figure S1b**-**c**). Spinal cord lesions from aCB_1_-KO mice at the chronic stage also showed reduced levels of SMI32 immunoreactivity, indicative of preserved neuroaxonal integrity, that were encompassed by an attenuated presence of microglia/macrophages and reactive astrocytes as determined by double immunolabelling for MBP and GFAP or Iba1 (**Figures S1d**-**f**). Thus, aCB_1_R exacerbates autoimmune inflammation and associated clinical symptomatology during EAE progression.

### CB_1_ receptor deletion prevents astrocyte dysfunction during EAE

To study the role of CB_1_R in the phenotypic transformation of astroglial cells during EAE we assessed the expression of molecules related to the acquisition of astrocyte pathogenic properties (43–45) in spinal cord tissue. Astrocyte-specific CB_1_R null mice showed reduced expression levels of several genes associated with the conversion of astroglial cells to disease-promoting phenotypes at acute EAE disease as determined by real-time qPCR analysis of spinal cord lysates (**Figure 1d**). We next examined inflammatory spinal cord lesions for the presence of complement component 3 (C3) as marker of pathogenic astrocytes in MS and EAE (44–46). Immunofluorescence staining of GFAP and C3 indicated reduced expression of both proteins in demyelinating spinal cord lesions from aCB_1_-KO mice (**Figure S2a**). Colocalization analysis evidenced lower C3 levels in astrocytic profiles (**Figure 1e**) associated to reduced numbers of microglia/macrophages (1863.5 ± 280 aCB_1_-KO *versus* 1342.8 ± 302 aCB_1_-WT; *p* = 0.014, Student’s *t* test). These results show that aCB_1_R promotes the astrocyte transformation into pathogenic phenotypes during EAE.

We next sought for possible differences between genotypes regarding astrocyte functional properties. Reactive astrocytes in the somatosensory cortex display impaired calcium responses that correlate to the severity of clinical symptomatology during EAE time-course (29, 46). Thus, we reasoned that astrocyte-specific CB_1_R deletion might attenuate glial reactivity and preserve astrocyte network function in this brain area. Fiber photometry analysis of cortical astrocytes Ca^2+^ activity in freely behaving aCB_1_-WT mice during EAE time-course (**Figure 1f**) showed reductions in the amplitude of sensory-evoked calcium signals at acute disease as compared to non-immunized mice (**Figure 1g**, *left panel*). This result resembles our recent observations on astrocyte calcium deregulation in this rodent model of MS using non-transgenic mice (46). Remarkably, astrocyte calcium responses to sensory stimulation were not significantly reduced in the brain cortex of aCB_1_-KO mice during the time-course of EAE (**Figure 1g**, *right panel*), which suggest an attenuation of disease-associated astrocyte hypo-responsiveness at the calcium signaling level in mutant mice. Cortical GFAP immunoreactivity was reduced in the aCB_1_-KO group when compared to aCB_1_-WT mice without reaching statistical significance (*p* = 0.0806; unpaired *t* test) (**Figure S2b**). However, histological examination showed general and astrocyte-specific reductions in C3 expression within deep cortical layers of aCB_1_-KO mice (**Figure 1h**; **Figure S2a**). Furthermore, immunofluorescence staining of CD3 revealed that the presence of cortical inflammatory lesions was also significantly reduced in aCB_1_-KO animals (**Figure 1i**). Thus, aCB_1_R deletion attenuates cortical inflammation and astrocyte network dysfunction at acute EAE disease.

### Mice with astrocytic CB_1_ receptor inactivation display intact oligodendrocyte populations in the EAE and LPC models

Oligodendrocyte differentiation prevents axonal loss and attenuates clinical symptomatology in the EAE model thus pinpointing to functional remyelination as potential mechanism of disease attenuation during autoimmune inflammation (47–49). Neurotoxic astrocytes release factors that promote oligodendrocyte apoptosis and delay lineage progression, leading to reduced remyelination and subsequent neuronal death (8, 45). Based on these evidence, we hypothesized that the protective phenotype in aCB_1_-KO mice in terms of astrocyte reactivity and demyelination extent may be related to the engagement of repair mechanisms during disease time-course. To determine whether aCB_1_R impede oligodendrocyte-differentiation promoting effects of astroglial cells as mechanism of clinical exacerbation, we immunostained spinal cord sections from aCB_1_-WT and aCB_1_-KO mice for the oligodendrocyte lineage marker Olig2 in combination with CC1 and NG2 to identify myelinating oligodendrocytes and OPCs, respectively. Astrocyte-specific CB_1_R mutants at acute EAE disease displayed unaltered numbers of Olig2^+^ oligodendrocyte lineage cells, CC1^+^/Olig2^+^ oligodendrocytes and NG2^+^/Olig2^+^ OPCs as compared to littermate controls, both in the demyelinating lesions and in the surrounding perilesion areas (**Figures 2a-d**). Consistently, the percentages of CC1^+^/Olig2^+^ mature oligodendrocytes and NG2^+^/Olig2^+^ OPCs in the plaques and periplaques were similar between aCB_1_-WT and aCB_1_-KO mice (**Figure 2e**). Thus, astrocyte CB_1_R does not modulate oligodendrocyte populations at acute EAE disease. Consistently, gene expression analysis of oligodendrocyte/myelin genes (*Olig2*, *Pdgfra, Mbp*, *Mog*) and factors that promote oligodendrocyte differentiation and (re)myelination (*Bdnf*, *Cntf*,*Ifg1*, *Ntf3*, *Pdgfa*, *Tgfb1*) did not show significant differences between genotypes (**Figure 2f**). Collectively, these results suggest that aCB_1_R do not hinder myelin repair as mechanism of clinical deterioration during EAE.

**Figure 2.**
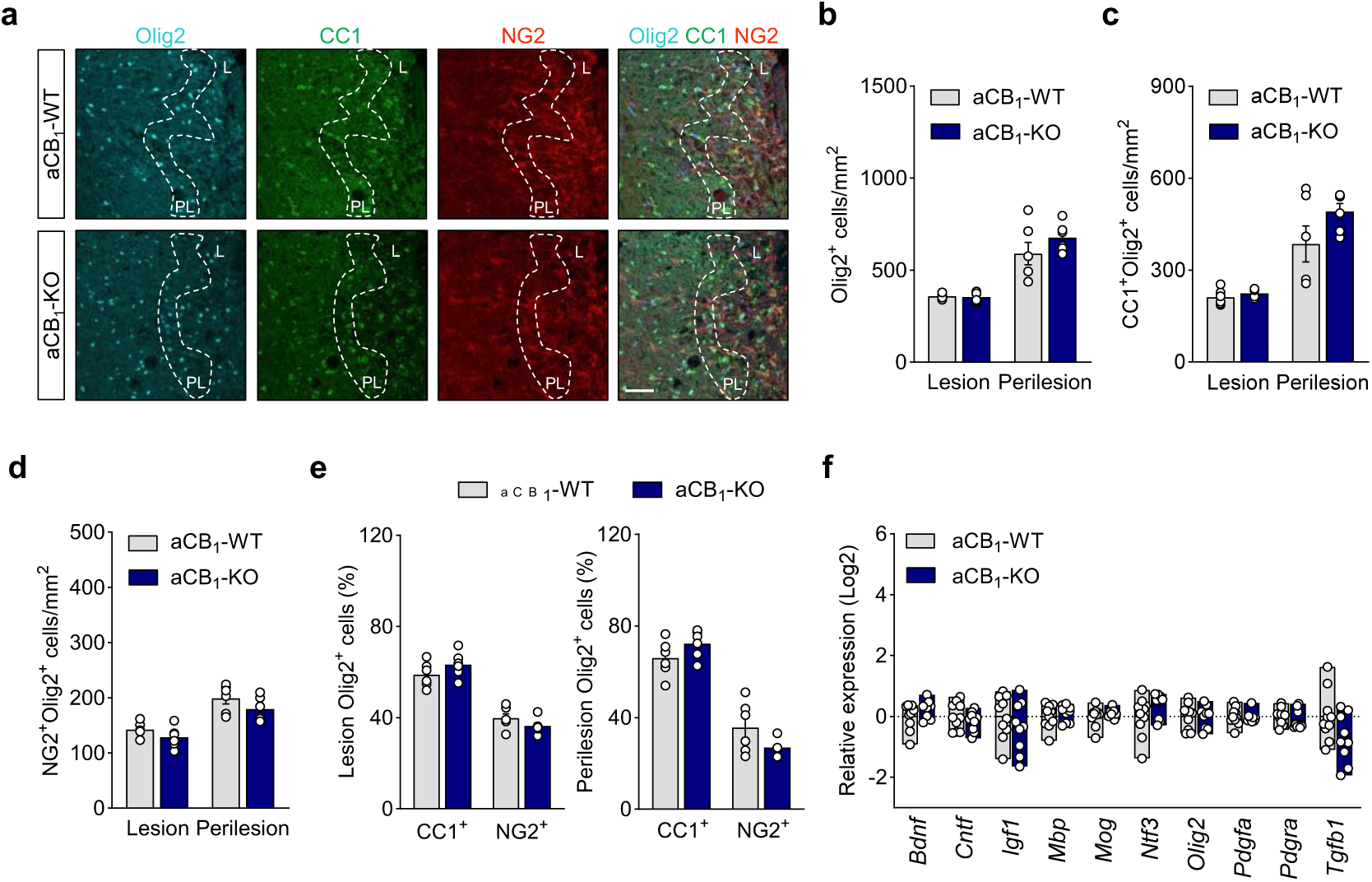
Astrocyte CB_1_R does not modulate oligodendrocyte populations in EAE inflammatory lesions. (**a**) Representative images of spinal cord lesions from aCB_1_-KO and aCB_1_-WT mice at acute EAE (25 dpi) immunostained for Olig2, CC1 and NG2 as markers of oligodendrocyte lineage cells, mature oligodendrocytes and oligodendrocyte precursor cells, respectively, show low cell densities in the lesion (L) and enrichment in the perilesion area (PL, dashed lines). Scale bar = 50 µm. (**b**-**d**) Quantitative analyses show similar densities of oligodendrocyte populations in the plaque and periplaque of aCB_1_-KO and aCB_1_-WT mice (*n* = 6 mice). (**e**) Percentages of oligodendrocytes (CC1^+^) and OPCs (NG2^+^) in the Olig2-positive population indicate equal proportions of mature cells in the lesion and perilesion of aCB_1_-KO and control littermates. (**f**) Gene expression analysis of oligodendrocyte/myelin markers and pro-myelinating molecules in spinal cord tissue from aCB_1_-KO and aCB_1_-WT mice at acute disease (22 dpi) (*n* = 9-10 mice).

Mechanistic studies of remyelination in EAE mice are challenging as autoimmune inflammation produces concomitant demyelination, axonal damage and myelin repair (50). To gain further insights on the role of aCB_1_R during remyelination *in vivo* we used a toxin-induced model in which focal lesions generated by localized injection of LPC in spinal cord white matter are followed by spontaneous remyelination (51, 52). OPCs are recruited into the demyelinated lesion and differentiate to mature oligodendrocytes during the second week post-lesion thus providing a defined time window to study changes in the rate of remyelination. We analyzed oligodendrocyte populations in LPC lesions from aCB_1_-WT and aCB_1_-KO mice at 14 days post-lesion (dpl) corresponding to the peak of endogenous OPC differentiation and the stage of the remyelination phase (**Figure S3a**). The demyelination extent of LPC lesions, assessed by co-immunostaining against MBP, was similar between phenotypes (**Figure S3b**). Quantification of double-labelled CC1^+^/Olig2^+^ oligodendrocytes and CC1^-^/Olig2^+^ oligodendrocyte precursors in LPC lesions revealed no variations between aCB_1_-KO and aCB_1_-WT mice (**Figures S3c**-**e**). Consistently, the numbers of PDGFR^+^ cells in LPC lesions were similar between aCB_1_-KO and aCB_1_-WT mice (**Figure S3f**). We also investigated potential changes in the population of the oligodendrocytes expressing brain-enriched myelin-associated protein 1 (BCAS1), which has been identified as a population of immature cells actively involved in (re)myelination (53, 54). Immunohistochemistry of LPC lesions for BCAS1^+^ revealed significant numbers of active oligodendrocytes in the lesion area of both aCB_1_-KO and aCB_1_-WT mice that were similar between genotypes (**Figure S3g**). Thus, aCB_1_R does not modulate spontaneous OPC differentiation following toxic oligodendrocyte loss. We next assessed LPC lesions from aCB_1_-KO and aCB_1_-WT mice for the presence of inflammatory cells. GFAP expression was not significantly different between groups, suggesting that LPC injections activate astrogliosis to a similar extent in aCB_1_-KO and aCB_1_-WT mice (**Figure S3h**). The presence of Iba1^+^ microglia/macrophages and CD45 infiltrating inflammatory cells was similar between genotypes (**Figures S3i**-**j**). These combined results suggest that aCB_1_R do not interfere with myelin repair processes in inflammatory and toxin induced demyelinating contexts and point to alternative mechanisms for clinical exacerbation during autoimmune inflammation.

### Reduced humoral and cellular infiltration in EAE lesions from astrocyte-specific CB_1_ receptor null mice

Perivascular astrocyte processes are enriched in CB_1_R that modulate BBB permeability during stress-induced inflammation (55–58). Thus, we wondered whether changes in the BBB properties contribute to the protective phenotype of astrocyte-specific CB_1_ null mice during EAE. To address this question, we initially measured parenchymal entry of humoral factors and immune cells as readouts of BBB opening at acute EAE disease (37, 59, 60). The areas immunopositive for the serum proteins fibrinogen and IgG as markers of humoral factor extravasation were markedly reduced in spinal cord lesions from aCB_1_-KO mice (**Figure 3a**). Moreover, spinal cord tissues from aCB_1_-KO mice contained significantly lower numbers of CD45^+^ leukocytes (**Figure 3b**) and densities of CD3^+^, B220^+^ and Ly6G^+^ leukocyte subsets (**Figure 3c**) within EAE lesions or perilesion areas, as measured using histopathology. Thus, aCB_1_-KO mice display preserved BBB function during autoimmune inflammation.

**Figure 3.**
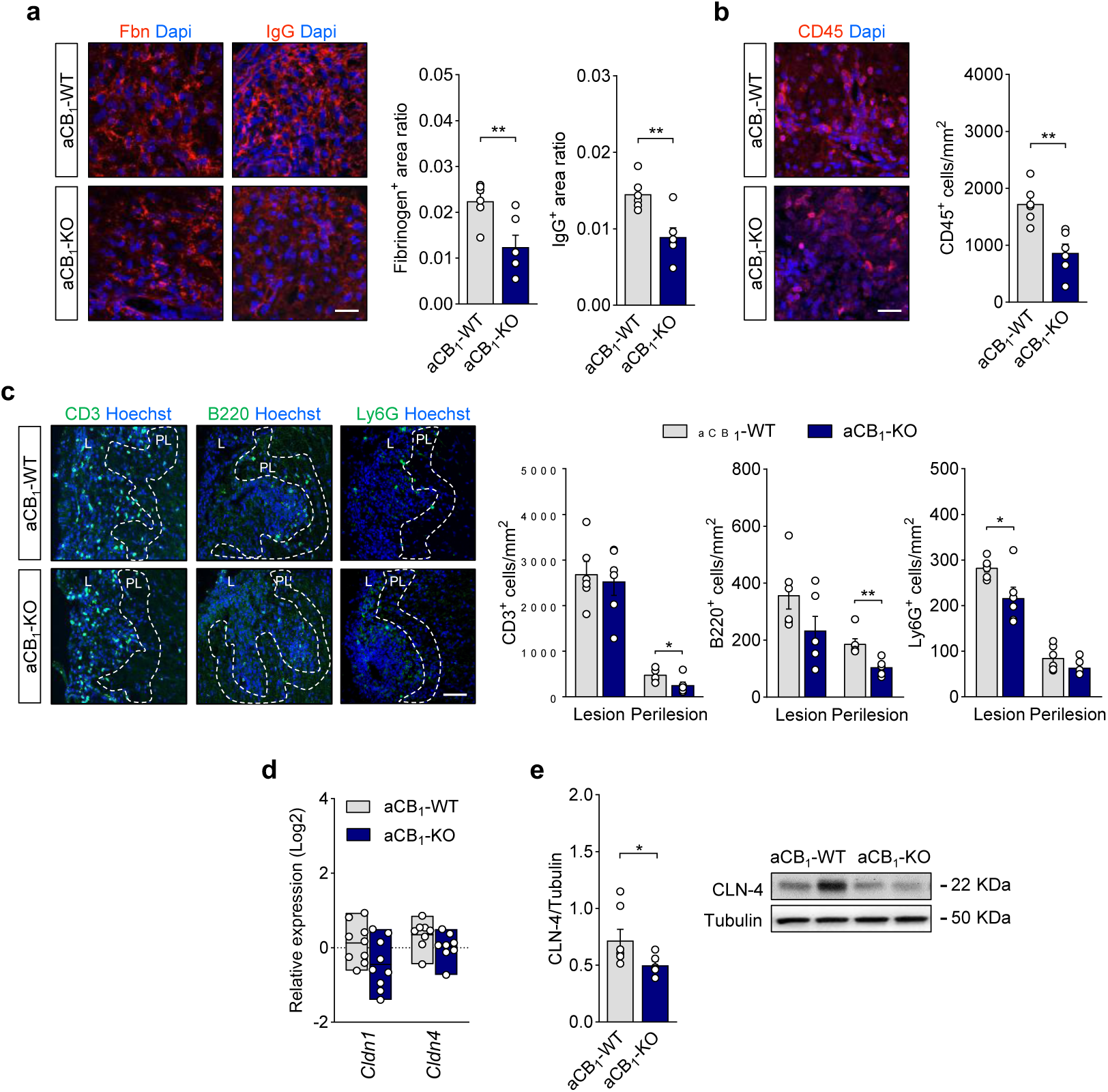
Restricted humoral and immune cell infiltration in EAE inflammatory lesions of aCB_1_R null mice. (**a**) Representative confocal z-stack projections of spinal cord lesions from aCB_1_-KO and control aCB_1_-WT mice at acute EAE (25 dpi) immunostained for fibrinogen (Fbn) and IgG. Morphometry of parenchymal immunoreactivity shows reduced extravasation in aCB_1_-KO mice as compared to the aCB_1_-WT group (*n* = 6 mice). Scale bar = 25 µm. (**b**) Representative micrograph and quantification of CD45^+^ infiltrating inflammatory cells in demyelinating lesions of aCB_1_-KO and aCB_1_-WT mice (*n* = 6 mice). Scale bar = 25 µm. (**c**) Confocal images and quantitative analysis of CD3^+^ T cells, B220^+^ B cells and Ly6G^+^ neutrophils in lesion (L) and perilesion (PL, dashed lines) spinal cord tissue from aCB_1_-KO mice as compared to aCB_1_-WT animals at acute EAE disease (*n* = 6 mice). Scale bar = 50 µm. (**d**) Gene expression analysis of the astrocyte TJ proteins CLN-1 andCLN-4 in spinal cord tissue from aCB_1_-KO and aCB_1_-WT mice at 22 dpi (*n* = 9-10 mice). (**e**) Immunoblots and densitometry analysis of CLN-4 expression in spinal cord lysates from aCB_1_-KO at 25 dpi compared to the aCB_1_-WT group (*n* = 6-7 mice). \**p* < 0.05, \*\**p* < 0.01 and \*\*\**p* < 0.001, unpaired *t*-test or Mann-Whitney test.

Astrocytes promote the formation of demyelinating lesions in EAE and MS by mechanisms that lead to the repression of endothelial tight junction (TJ) proteins and drive BBB permeability (59, 60). Concomitantly, astroglial cells activated in response to neuroinflammation upregulate claudin 1 (CLN-1) and claudin 4 (CLN-4) to form protective astrocytic TJ complexes that control humoral and cellular transit (61). Expression of *Cldn1* and *Cldn4* in spinal cord tissue was not significantly modulated in the aCB_1_-KO as compared to aCB_1_-WT mice (**Figure 3d**). However, western blot analysis of spinal cord lysates showed reduced expression levels of the main astrocytic TJ protein CLN-4 (61) (**Figure 3e**), which are consistent with the attenuated inflammatory phenotype of aCB_1_-KO mice (**Figures 3b-c**). To investigate whether aCB_1_R modulates the endothelial BBB during autoimmune inflammation we analyzed the expression of vascular endothelial cell markers and TJ associated proteins in spinal cord tissues from EAE mice. Astrocyte-specific CB_1_R null mice displayed unaltered levels of the vascular endothelial membrane proteins podocalyxin (PODXL) and laminin, as measured using immunoblotting and confirmed by immunohistochemistry of EAE lesions (**Figures S4a**-**b**). These observations suggest that aCB_1_R-KO mice do not display abnormal angiogenesis at acute EAE disease as compared to wild-type animals. The expression of the endothelial TJ components cadherin 5 (CDH-5), zonula occludens (ZO-1) and claudin 5 (CLN-5) was also similar between aCB_1_-WT and aCB_1_-KO mice (**Figures S4c**-**d**). Thus, aCB_1_R null mice display microvascular endothelial cells similar to wild-type controls at established EAE disease despite the attenuated humoral and leukocyte infiltration in lesion and perilesion sites.

### Mice with astrocyte CB_1_ inactivation show reduced expression of immune and vascular effector molecules in EAE lesions

Reactive astrocytes contribute to the pathogenesis of CNS inflammatory lesions thought the release of intercellular molecules that enable circulating inflammation to reach the brain and spinal cord parenchyma. Astrocyte effector molecules that mediate autoimmune inflammation include the chemokines CCL2, CCL5 and CXCL2 as recruiters of perivascular leukocytes (8, 10), adhesion molecules such as ICAM-1 and VCAM-1 aberrantly expressed by astroglial cells (62–65), and angiogenic factors, mainly VEGF-A, that signal to the vascular endothelium and promote permeability (37, 59, 60). To gain further insights on the mechanistic implications of aCB_1_R during autoimmune inflammation we addressed the expression of immune and vascular effector molecules at acute EAE disease. The expression levels of *Ccl2*, *Ccl5* and *Cxcl2* determined by RT-qPCR in spinal cord lysates of EAE mice were highly heterogeneous and not significantly modulated in the aCB_1_-KO group (**Figure 4a**). Conversely, gene expression analysis highlighted significantly lower levels of *Icam1* and *Vcam1* that were confirmed by immunoblotting and confocal imaging of spinal cord tissue (**Figures 4a-b** and **S5a**-**b**). Moreover, double immunostaining for GFAP and ICAM-1 or VCAM-1 showed reduced expression of both adhesion molecules in astrocytic profiles within EAE lesions and perilesion areas of aCB_1_-KO mice (**Figure 4c**). Of note, VEGF-A expression was drastically downregulated in spinal cord tissue and reactive astrocyte profiles of aCB_1_-KO mice as compared to the aCB_1_-WT group (**Figures 4c** and **S5a**-**b**). Thus, the inhibition of humoral and lymphocyte infiltration in white matter lesions of mice with aCB_1_R deletion is accompanied by a restricted astrocyte production of adhesion molecules and VEGF-A during autoimmune inflammation.

**Figure 4.**
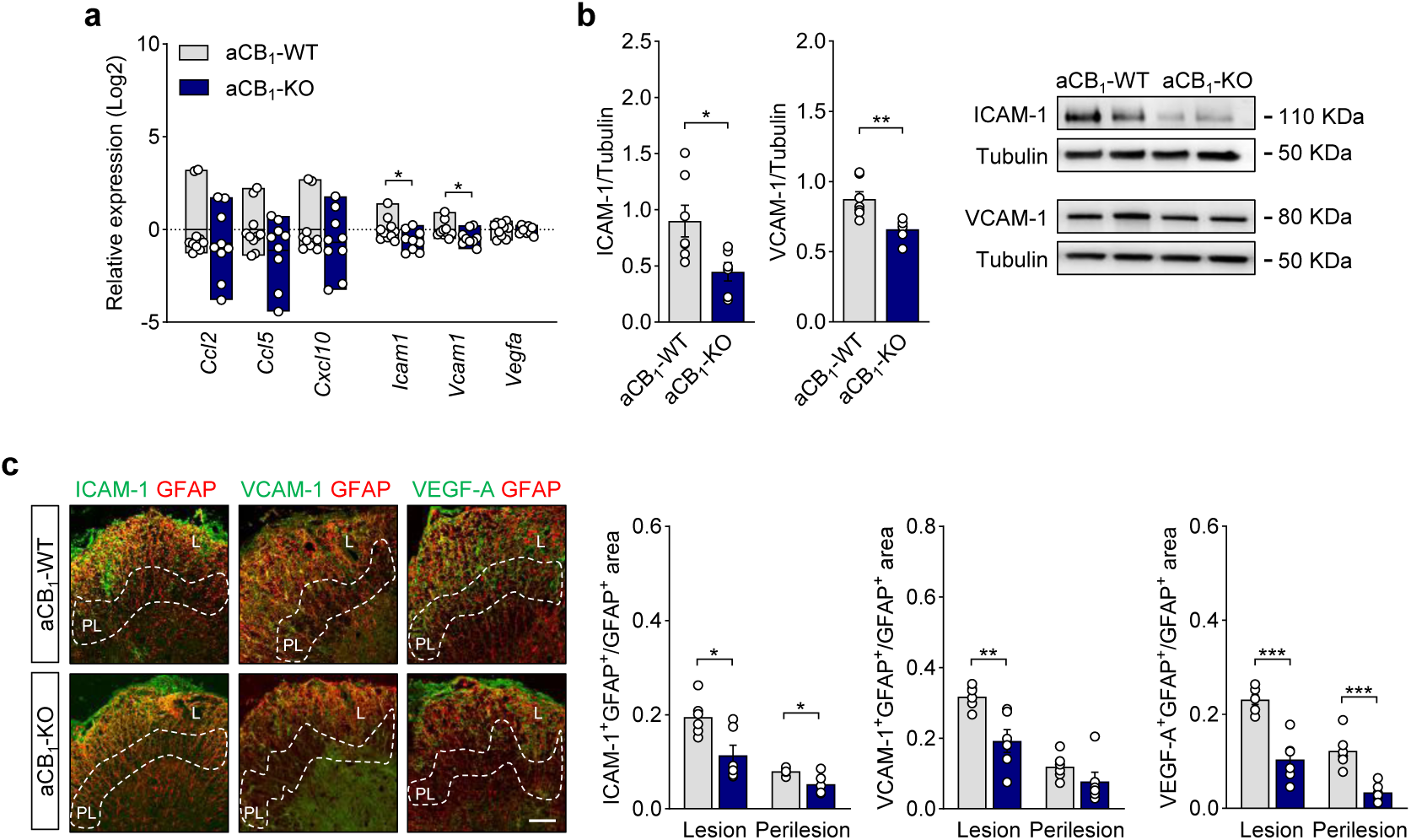
Astrocyte-specific CB_1_R null mice show reduced expression of immune and vascular effector molecules in EAE lesions. (**a**) Relative gene expression of permeability effector molecules in spinal cord tissue from aCB_1_-KO mice as compared to control aCB_1_-WT animals at acute disease EAE (*n* = 9-10 mice). (**b**) Immunoblotting for the adhesion molecules ICAM-1 and VCAM-1 shows reduced expression levels in aCB_1_-KO spinal cords (n = 6-7). (**c**) Confocal z-stack projections of spinal cord sections immunostained for GFAP and ICAM-1, VCAM-1 or VEGF-A and colocalization analysis highlight attenuated expression of permeability molecules in astroglial profiles at lesion (L) and/or perilesion (PL, dashed lines) areas from aCB_1_-KO animals (*n* = 6 mice). Scale bar = 100 µm. \**p* < 0.05, \*\**p* < 0.01 and \*\*\**p* < 0.001, unpaired *t*-test or Mann-Whitney test.

### Diminished inflammatory neuropathology of astrocyte CB_1_ receptor null mice in the VEGF-A model of BBB breakdown

Collectively, the above findings suggest that aCB_1_R facilitate BBB permeability to promote inflammatory lesion formation during autoimmune inflammation. To further investigate potential links between astrocyte CB_1_R and BBB breakdown *in vivo*, we analyzed the effects of VEGF-A in astrocyte-specific CB_1_R null mice. We stereotactically delivered mouse VEGF-A_165_ into the left cerebral cortex of aCB_1_-KO mice and littermate aCB_1_-WT animals and measured changes in barrier function at 2 days post-lesion (**Figure 5a**). Compatible with published data (37, 60), we observed BBB breakdown measured by fibrinogen and IgG immunoreactivity in VEGF-A_165_-injected areas (**Figure 5b**). The endothelial TJ markers CDH-5 and ZO-1 appeared patchy and discontinuous in both aCB_1_-WT and a-CB_1_-KO mice and these changes were accompanied by parenchymal accumulation of CD45^+^ cells (**Figure 5b**). Importantly, BBB disruption, as measured by serum protein and immune cell extravasation, was attenuated in aCB_1_-KO mice despite unaltered expression levels of CHD-5, ZO-1 and the endothelial cell maker PODXL (**Figures 5c-d**). Reminiscent of our observations in the EAE lesions, aCB_1_R null mice showed reduced astrocyte reactivity (**Figure 5e**), and lower levels of the astrocyte TJ protein CLN-4 in areas of BBB disruption but not of CLN-1(**Figure 5f**). These studies show that aCB_1_R engages mechanisms downstream of VEGF-A that promote BBB leakiness and allows for the infiltration of peripheral immune during inflammatory lesion formation.

**Figure 5.**
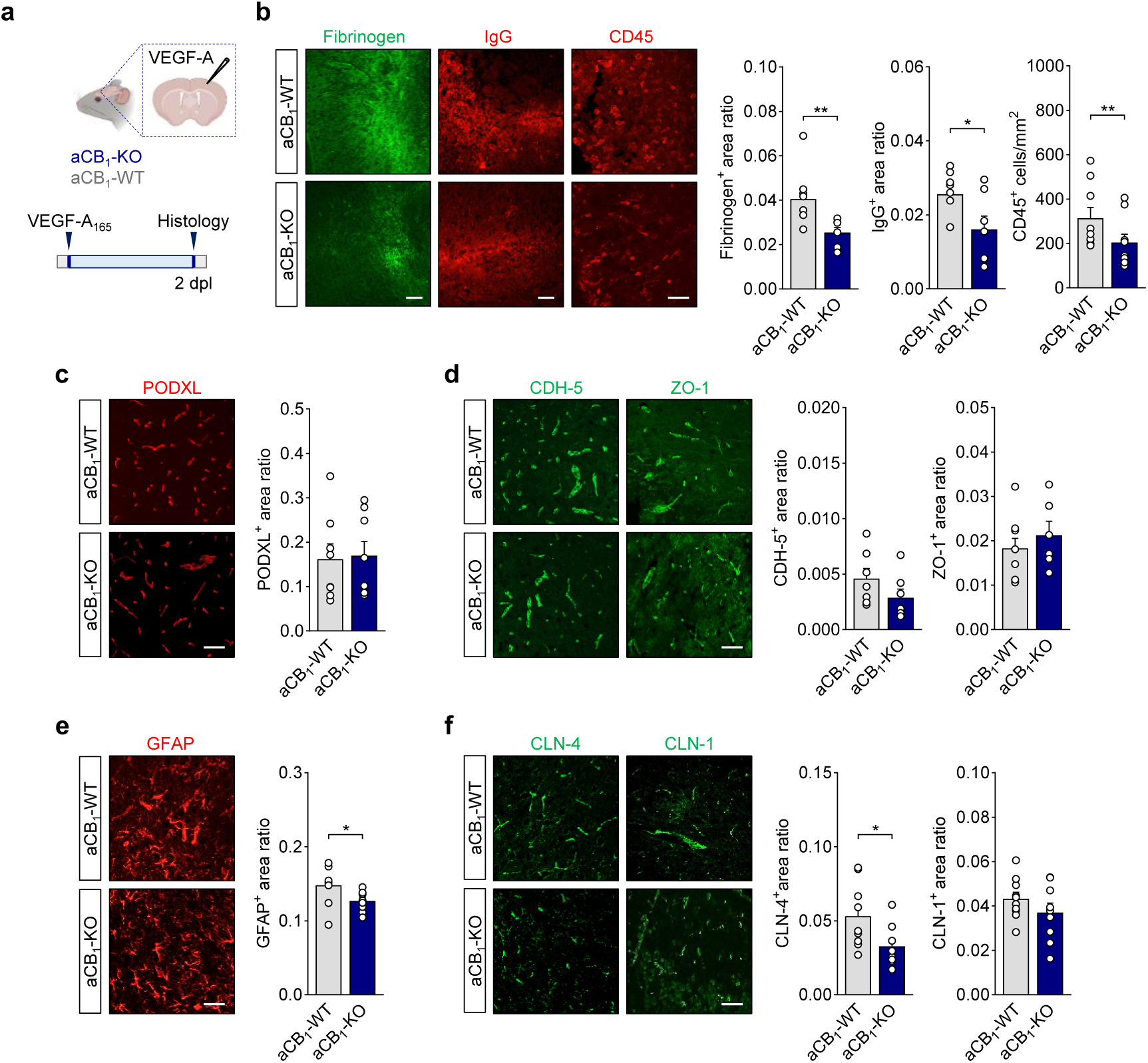
Deletion of astrocyte CB_1_R reduces cortical BBB leakage, astrocytic tight-junction deregulation and leukocyte infiltration induced by VEGF-A. (**a**) Experimental design for the analysis of the BBB breakdown following cortical microinjection of murine VEGF-A_165_ in aCB_1_-KO and aCB_1_-WT mice. Cerebral cortices from animals injected in 2 independent experiments were processed for histopathology at 2 days post-lesion (dpl). (**b**) Representative cortical tissue sections immunostained for fibrinogen, IgG and CD45 and quantitative analysis show reduced extravasation of serum proteins and attenuated leukocyte infiltration in the aCB_1_-KO group. (*n* = 9 mice). Scale bars = 100 µm (fibrinogen, IgG) and 50 µm (CD45). (**c**, **d**) Confocal micrographs and morphometry of PODXL (**c**) and endothelial TJ proteins CDH-5 and ZO-1 (**d**) in aCB_1_-KO and aCB_1_-WT mice. Scale bar = 50 µM. (**e**, **f**) Attenuated astrocyte reactivity measured by immunostaining for GFAP (**e**) and levels of astrocyte TJ proteins CLN-4 and CLN-4 (**f**) following microinjection of VEGF-A_165_ in aCB_1_-KO and aCB_1_-WT mice. Scale bar = 50 µm. \**p* < 0.05 and \*\**p* < 0.01, unpaired *t*-test or Mann-Whitney test.

## Discussion

Studies conducted during the past decades have demonstrated that CB_1_R signaling restricts clinical disability in MS (16, 21). Potential mechanisms for (endo)cannabinoid-mediated protection mediated by CB_1_R in MS patients include neuroprotective and remyelination promoting effects, as defined by analyzing the phenotype of constitutive CB_1_R-KO mice (22) and transgenic mice lacking CB_1_R on neurons (20) and OPCs (25) in preclinical disease models. However, the specific roles of CB_1_R expressed by astrocytes in MS have remained largely neglected despite the critical involvement of these cells in disease initiation and progression (8, 10), and the emerging implications of the astrocyte CB_1_R pools in endocannabinoid-related modulation of brain functions (66, 67). This study is the first to our knowledge that highlights the significance of CB_1_R endocannabinoid signaling mediated by astrocytes in the generation of clinical deficits in CNS inflammatory disease. Upon characterizing the phenotype of mice lacking aCB_1_R in models of acute and chronic CNS damage, we depict an unprecedented deleterious role of CB_1_R in astroglial cells during inflammatory lesion formation linked to the modulation of BBB permeability. On mechanistic grounds, we show that aCB_1_R deletion restricts humoral and cellular leakage associated to BBB disruption, at least in part, by engaging cellular processes downstream VEGF-A signaling. These observations uncover a novel mechanism underlying astrocyte-related control of the BBB in inflammatory CNS disease and shed light on the roles of endocannabinoids and CB_1_R in MS pathogeny.

In this study, we show protective effects of astrocytic CB_1_R inactivation during the time-course of EAE at the clinical and neuropathological level. Attenuation of neurological disability occurred in concert with changes in astrocyte reactivity and expression of molecules related to the acquisition of disease promoting functions in MS patients and preclinical disease models (43, 45). These observations are consistent with a scenario in which astrocytic CB_1_R signaling facilitates the emergence of neurotoxic phenotypes that impede astrocyte-oligodendrocyte interactions underlying myelin repair as potential mechanism of disease exacerbation (8). Studies specifically designed to investigate the involvement of cell autonomous astrocyte responses mediated by CB_1_R in remyelination are currently lacking. Thus, we addressed this possibility by histologically targeting oligodendrocyte populations in lesion and perilesion areas of aCB_1_-KO mice at acute EAE disease. This analysis did not evidence significant modulatory effects encompassing the attenuation of inflammation and myelin pathology that resulted from deletion of astrocytic CB_1_R. Consistently, analysis of remyelinating spinal cord lesions induced by microinjection of LPC did not highlight oligodendrocyte differentiation promoting effects in astrocyte-specific CB_1_R null mice. Although the possibility that endocannabinoids target CB_1_R in astroglial cells to modulate remyelination *in vivo* cannot be fully disregarded in the absence of a detailed myelin ultrastructure analysis during the time-course of myelin repair, these results suggest that astrocytic CB_1_R do not impede remyelination as potential mechanism of clinical exacerbation in demyelinating disease.

Our present data support the hypothesis that astrocyte responses to CB_1_R signaling modulate the formation of CNS lesions as principal mechanism of disease exacerbation. This possibility sits well with the fundamental roles of astroglial cells in regulating the BBB breakdown during CNS inflammation (8, 10) and with reports of enriched CB_1_R expression in perivascular astrocytic profiles (55, 57, 58). Indeed, the clinical benefits of astrocyte-specific CB_1_R inactivation during EAE are observed at the onset of neurological symptomatology and occur in concert with reduced inflammatory lesion load at established disease, both in cortical and spinal cord tissue. These observations prompted us to investigate the phenotype of aCB_1_-KO mice in terms of BBB disruption during EAE. Here, as expected if CB_1_R in astrocytes would promote BBB permeability defects allowing circulating inflammation to enter the CNS parenchyma, aCB_1_-KO mice exhibited lower levels of humoral mediators and infiltrating leukocytes in inflammatory demyelinating EAE lesions and surrounding perilesion areas. Notably, these changes were associated with reduced expression levels of the adhesion molecules ICAM-1 and VCAM-1 by reactive astrocytes in EAE lesions. These findings may explain, at least in part, the restricted presence of infiltrating immune cells observed in mice with astrocyte-specific CB_1_R deletion, according to the established roles of both adhesion molecules in mediating leukocyte movement across the BBB during CNS inflammatory disease (62–65). Reactive astrocytes in EAE lesions from aCB_1_-KO mice also exhibited lower levels of the angiogenic and pro-permeability factor VEGF-A, identified as a key driver of astrocyte-mediated BBB disruption, leukocyte infiltration and neuropathology during CNS inflammation (59, 60). Reductions in astrocyte VEGF-A expression might directly limit leukocyte entry as primary mechanism underlying reduced lesion load in astrocyte-specific CB_1_R null mice. Concomitantly, lower levels of adhesion molecules or other potential astrocyte mediators may further restrict BBB permeability as evolving inflammation drives astrocyte VEGF-A expression (68). Using the focal, directly induced VEGF-A model in transgenic mice allowed us to selectively examine the role of CB_1_R-related astrocyte responses on BBB breakdown and lymphocyte entry downstream this vascular effect molecule. Critically, mice with astrocyte-specific CB_1_R deletion showed milder humoral and cellular extravasation induced by VEGF-A, thus mirroring the restricted inflammatory phenotype observed in EAE lesions. Altogether, these findings suggest that CB_1_R signaling in astrocytes facilitates the formation of CNS inflammatory lesions by targeting effector mechanisms engaged by VEGF-A that promote BBB disruption.

Our combined observations reflect a general protective astrocytic phenotype in terms of BBB permeability resulting from CB_1_R deletion in astroglial cells. Notably, these changes occurred in concert with reduced levels of the astrocyte TJ protein CLN-4, whose induction has been associated to the emergence of a protective astrocyte barrier that limits the access of peripheral inflammation to the CNS parenchyma (61). These seemingly contradictory findings can be explained on the basis that the evolving inflammation in lesions sites, which we find to be significantly reduced in mice lacking aCB_1_R, is a driving force for the induction of junctional adhesion proteins at the glia limitants (61). A striking observation in this study, however, is that attenuated parenchymal leakage of circulating inflammation in lesions from astrocyte-specific CB_1_R was not encompassed by significant endothelium abnormalities, as expected according to reports that BBB breakdown results from disruptions of endothelial junctional proteins such as CLN-5 (37, 59, 60). Nevertheless, our present findings do not exclude the possibility that astrocyte-specific CB_1_R null mice exhibit preserved endothelial barrier functions at initial stages of BBB dysfunction in the EAE and VEGF-A models, as compared to non-transgenic animals. Disruption of the BBB precedes immune cell infiltration during CNS lesion formation (37, 69) and it seems plausible that early-onset differences in endothelial permeability between genotypes are masked at later states by the evolving inflammatory neuropathology. Future studies targeting the time-course of CNS lesion formation *in vivo* combined with clinically relevant *in vitro* models of BBB dysfunction may uncover the mechanistic basis of astrocyte signaling to endothelial cells under the control of endocannabinoids and astrocytic CB_1_R pools that lead to clinical exacerbation during demyelinating disease.

Endocannabinoids and their exogenous counterparts dampen CNS inflammation in multiple disease paradigms (14, 16). However, there is a paucity of studies that investigate the signaling events engaged by these lipid mediators at the BBB permeability level (56, 70, 71). In this context, available evidence supports a protective role for CB_1_R expressed by endothelial cells in attenuating leukocyte transmigration induced by Theiler’s virus infection (72). More recently, perivascular astrocyte CB_1_R has been postulated to promote stress resilience in mice through the preservation of BBB functions (56). In this landmark study, viral-mediated astrocyte overexpression of CB_1_R mitigated inflammatory responses and morphological changes in the chronic social defeat stress (CSDS) mouse model of depression. These observations are in apparent contrast with our present findings that astrocyte CB_1_R deletion preserves BBB function and restrains circulating inflammation to enter the brain parenchyma. It should be noted, however, that stress-induced inflammation and autoimmune demyelination are intrinsically different disease contexts that involve dissimilar etiopathological mechanisms. Furthermore, the protective phenotype of astrocyte CB_1_R at the vascular level reported by Dudek and collaborators was ascribed to astrocyte populations within the nucleus accumbens shell in male mice resilient to CSDS. Conversely, this study, that we performed in female mice, suggests more generalized mechanisms underlying the control of BBB permeability by astrocytic CB_1_R during CNS lesion formation, as aCB_1_-KO mice showed attenuated spinal cord white matter and cortical grey matter inflammatory pathology in complementary disease models. Remarkably, CSDS and viral-induced downregulation of endothelial *Cln5* increase astrocyte CB_1_R and endocannabinoid levels *in vivo* while acute inflammatory challenges with IL-6 upregulate the expression levels of *Cnr1* in culture systems (56). These recent observations suggest that BBB disruption facilitates astrocytic CB_1_R signaling though inflammation-related mechanisms. In this scenario, it seems plausible that distinct inflammatory environments engage endocannabinoid signaling through specific astrocyte CB_1_R receptor pools at the plasma membranes and/or mitochondrial compartments, leading to context specific modulation of BBB functions. Mechanistic considerations notwithstanding, our study adds to the growing body of evidence that support essential roles for the astrocytic populations of CB_1_R in regulating the astrocyte-endothelial interface in neuroinflammatory disorders.

The exact cascade of events that modulate BBB permeability downstream astrocytic CB_1_R signaling during CNS lesion formation remains a matter of future research efforts in the context of autoimmune inflammation. According to the existing literature, early-onset activation of astrocytic CB_1_R may potentially preserve BBB function by attenuating astrocyte reactivity (8, 10). This possibility is supported by a series of *in vivo* ablation studies showing that early onset inhibition of reactive astrogliosis facilitates circulating inflammation to reach the CNS parenchyma and exacerbates clinical disease in the EAE model of MS (73–75). Some tentative additional support for this hypothesis comes from prior research showing that pharmacological blockade of endocannabinoid hydrolysis inhibits astrocyte reactivity while attenuating disease neuropathology and clinical severity in the EAE and TMEV models of neuroinflammation (76, 77). However, these *in vivo* studies did not target CB_1_R as potential underling mechanism and research specifically designed to address the role of CB_1_R in regulating the functions of reactive astrocytes is still scarce. In this context, it is worth mentioning that the protective phenotype of astrocyte-specific CB_1_R null mice in the EAE model that we report here is consistent with recent *in vivo* observations of decreased neuronal death following cerebral ischemia in mice (78).

To summarize, we propose that early, endogenous activation of astrocyte CB_1_R drives the pathogenicity of CNS inflammatory lesions by mechanistic interactions with VEGF-A signaling that promote BBB permeability. This study challenges the traditional neuroprotective role of endocannabinoids in MS and adds to the accumulating evidence that point to relevant roles of endocannabinoid signaling via astrocyte CB_1_R in the neurovascular adaptations that shape neuroinflammation.

## Data availability

The datasets generated during the current study are available from the corresponding authors on reasonable request.

## Abbreviations

aCB_1_R: Astrocyte cannabinoid type-1 receptor
BBB: Blood-brain barrier
CB_1_R: Cannabinoid type-1 receptor
CDH-5: Cadherin 5
cDNA: Complementary DNA
CLN-1: Claudin 1
CLN-4: Claudin 4
CLN-5: Claudin 5
CNS: Central nervous system
GFAP: Glial fibrillary acid protein
ICAM-1: Intercellular adhesion molecule 1
LED: Light-emitting diode
LFB: Luxol fast blue
LPC: lysophosphatidylcholine
MBP: Myelin basic protein
MOG: Myelin oligodendrocyte glycoprotein
MS: Multiple sclerosis
NDS: Normal donkey serum
NGS: Normal goat serum
OPC: Oligodendrocyte precursor cell
PBS: Phosphate buffered saline
PCR: Polymerase chain reaction
PODXL: Podocalyxin
RIPA: Radioimmunoprecipitation assay
RNA: Ribonucleic acid
RT: Room temperature
TBS: Tris buffer saline
THC: Δ^9^-tetrahydrocannabinol
VCAM-1: Vascular cell adhesion protein 1
VEGF-A: Vascular endothelial growth factor A
ZO-1: Zonula occludens

## Acknowledgements

We thank all members of the S Matós and G Marsicano’s lab for useful discussions and advice, as well as the personnel of the Animal Facilities of the University of the Basque Country and Neurocentre Magendie for mouse care.

## Author contributions

T.C., E.S.-M., A.B.-C., and A.U, performed and analyzed histological, immunoblot and gene expression experiments. A.M.-G. and R.S. performed fiber photometry analysis. U.S. and I.F-M. were involved in mouse breeding and performed pharmacological treatments. V.T. conceptualized LPC experiments, performed surgeries and was involved in data analysis and interpretation. C.C. performed cortical surgeries and provided guidance for data analysis and interpretation. C.M. participated in EAE experiments. A.M.B. was involved in the analysis of calcium imaging experiments. G.M. and S.M. conceptualized and supervised the study. A.M.B., T.C. and S.M. produced the figures and wrote the paper. All authors approved the final manuscript.

## References

1. Reich DS, Lucchinetti CF, Calabresi PA. Multiple sclerosis. N Engl J Med. 2018;378(2):169–80.

2. Mahad DH, Trapp BD, Lassmann H. Pathological mechanisms in progressive multiple sclerosis. Lancet Neurol. 2015;14(2):183–93.

3. Filippi M, Bar-Or A, Piehl F, Preziosa P, Solari A, Vukusic S, et al. Multiple sclerosis. Nat Rev Dis Primers. 2018;4(1):43.

4. Lazzarotto A, Hamzaoui M, Tonietto M, Dubessy AL, Khalil M, Pirpamer L, et al. Time is myelin: early cortical myelin repair prevents atrophy and clinical progression in multiple sclerosis. Brain. 2024;147(4):1331–43.

5. Lubetzki C, Zalc B, Williams A, Stadelmann C, Stankoff B. Remyelination in multiple sclerosis: from basic science to clinical translation. Lancet Neurol. 2020;19(8):678–88.

6. Escartin C, Galea E, Lakatos A, O’Callaghan JP, Petzold GC, Serrano-Pozo A, et al. Reactive astrocyte nomenclature, definitions, and future directions. Nat Neurosci. 2021;24(3):312–25.

7. Hasel P, Rose IVL, Sadick JS, Kim RD, Liddelow SA. Neuroinflammatory astrocyte subtypes in the mouse brain. Nat Neurosci. 2021;24(10):1475–87.

8. Linnerbauer M, Wheeler MA, Quintana FJ. Astrocyte crosstalk in CNS inflammation. Neuron. 2020;108(4):608–22.

9. Ponath G, Park C, Pitt D. The role of astrocytes in multiple sclerosis. Front Immunol. 2018;9:217.

10. Sofroniew MV. Astrocyte barriers to neurotoxic inflammation. Nat Rev Neurosci. 2015;16(5):249–63.

11. Gorter RP, Baron W. Recent insights into astrocytes as therapeutic targets for demyelinating diseases. Curr Opin Pharmacol. 2022;65:102261.

12. Molina-Gonzalez I, Holloway RK, Jiwaji Z, Dando O, Kent SA, Emelianova K, et al. Astrocyte-oligodendrocyte interaction regulates central nervous system regeneration. Nat Commun. 2023;14(1):3372.

13. Pertwee RG, Howlett AC, Abood ME, Alexander SP, Di Marzo V, Elphick MR, et al. International union of basic and clinical pharmacology. LXXIX. Cannabinoid receptors and their ligands: beyond CB₁ and CB₂. Pharmacol Rev. 2010;62(4):588–631.

14. Cristino L, Bisogno T, Di Marzo V. Cannabinoids and the expanded endocannabinoid system in neurological disorders. Nat Rev Neurol. 2020;16(1):9–29.

15. Lutz B. Neurobiology of cannabinoid receptor signaling. Dialogues Clin Neurosci. 2020;22(3):207–22.

16. Bernal-Chico A, Tepavcevic V, Manterola A, Utrilla C, Matute C, Mato S. Endocannabinoid signaling in brain diseases: Emerging relevance of glial cells. Glia. 2022;71(1):103–126.

17. de Lago E, Moreno-Martet M, Cabranes A, Ramos JA, Fernández-Ruiz J. Cannabinoids ameliorate disease progression in a model of multiple sclerosis in mice, acting preferentially through CB_1_ receptor-mediated anti-inflammatory effects. Neuropharmacology. 2012;62(7):2299–308.

18. Croxford JL, Miller SD. Immunoregulation of a viral model of multiple sclerosis using the synthetic cannabinoid R+WIN55,212. J Clin Invest. 2003;111(8):1231–40.

19. Arévalo-Martín A, Vela JM, Molina-Holgado E, Borrell J, Guaza C. Therapeutic action of cannabinoids in a murine model of multiple sclerosis. J Neurosci. 2003;23(7):2511–6.

20. Maresz K, Pryce G, Ponomarev ED, Marsicano G, Croxford JL, Shriver LP, et al. Direct suppression of CNS autoimmune inflammation via the cannabinoid receptor CB_1_ on neurons and CB_2_ on autoreactive T cells. Nat Med. 2007;13(4):492–7.

21. Chiurchiù V, van der Stelt M, Centonze D, Maccarrone M. The endocannabinoid system and its therapeutic exploitation in multiple sclerosis: Clues for other neuroinflammatory diseases. Prog Neurobiol. 2018;160:82–100.

22. Pryce G, Ahmed Z, Hankey DJ, Jackson SJ, Croxford JL, Pocock JM, et al. Cannabinoids inhibit neurodegeneration in models of multiple sclerosis. Brain. 2003;126(Pt 10):2191–202.

23. Rossi S, Furlan R, De Chiara V, Muzio L, Musella A, Motta C, et al. Cannabinoid CB_1_ receptors regulate neuronal TNF-α effects in experimental autoimmune encephalomyelitis. Brain Behav Immun. 2011;25(6):1242–8.

24. Croxford JL, Pryce G, Jackson SJ, Ledent C, Giovannoni G, Pertwee RG, et al. Cannabinoid-mediated neuroprotection, not immunosuppression, may be more relevant to multiple sclerosis. J Neuroimmunol. 2008;193(1-2):120–9.

25. Sánchez de la Torre A, Ezquerro-Herce S, Huerga-Gómez A, Sánchez-Martín E, Chara JC, Matute C, et al. CB₁ receptors in NG2 cells mediate cannabinoid-evoked functional myelin regeneration. Prog Neurobiol. 2024;243:102683.

26. Aguado T, Huerga-Gómez A, Sánchez-de la Torre A, Resel E, Chara JC, Matute C, et al. Δ⁹-Tetrahydrocannabinol promotes functional remyelination in the mouse brain. Br J Pharmacol. 2021;178(20):4176–92.

27. Covelo A, Eraso-Pichot A, Fernández-Moncada I, Serrat R, Marsicano G. CB_1_R-dependent regulation of astrocyte physiology and astrocyte-neuron interactions. Neuropharmacology. 2021;195:108678.

28. Eraso-Pichot A, Pouvreau S, Olivera-Pinto A, Gomez-Sotres P, Skupio U, Marsicano G. Endocannabinoid signaling in astrocytes. Glia. 2023;71(1):44–59.

29. Baraibar AM, Colomer T, Moreno-García A, Bernal-Chico A, Sánchez-Martín E, Utrilla C, et al. Autoimmune inflammation triggers aberrant astrocytic calcium signaling to impair synaptic plasticity. Brain Behav Immun. 2024;121:192–210.

30. Marsicano G, Araque A. Of glue and pot: Endocannabinoid signaling in glial cells. Glia. 2023;71(1):3–4.

31. Walton C, King R, Rechtman L, Kaye W, Leray E, Marrie RA, et al. Rising prevalence of multiple sclerosis worldwide: Insights from the Atlas of MS, third edition. Mult Scler. 2020;26(14):1816–21.

32. Marsicano G, Wotjak CT, Azad SC, Bisogno T, Rammes G, Cascio MG, et al. The endogenous cannabinoid system controls extinction of aversive memories. Nature. 2002;418(6897):530–4.

33. Hirrlinger PG, Scheller A, Braun C, Hirrlinger J, Kirchhoff F. Temporal control of gene recombination in astrocytes by transgenic expression of the tamoxifen-inducible DNA recombinase variant CreERT2. Glia. 2006;54(1):11–20.

34. Han J, Kesner P, Metna-Laurent M, Duan T, Xu L, Georges F, et al. Acute cannabinoids impair working memory through astroglial CB_1_ receptor modulation of hippocampal LTD. Cell. 2012;148(5):1039–50.

35. Robin LM, Oliveira da Cruz JF, Langlais VC, Martin-Fernandez M, Metna-Laurent M, Busquets-Garcia A, et al. Astroglial CB₁ receptors determine synaptic d-serine availability to enable recognition memory. Neuron. 2018;98(5):935–44.e5.

36. Tepavčević V, Kerninon C, Aigrot MS, Meppiel E, Mozafari S, Arnould-Laurent R, et al. Early netrin-1 expression impairs central nervous system remyelination. Ann Neurol. 2014;76(2):252–68.

37. Argaw AT, Gurfein BT, Zhang Y, Zameer A, John GR. VEGF-mediated disruption of endothelial CLN-5 promotes blood-brain barrier breakdown. Proc Natl Acad Sci U S A. 2009;106(6):1977–82.

38. Akerboom J, Carreras Calderón N, Tian L, Wabnig S, Prigge M, Tolö J, et al. Genetically encoded calcium indicators for multi-color neural activity imaging and combination with optogenetics. Front Mol Neurosci. 2013;6:2.

39. Ohkura M, Sasaki T, Sadakari J, Gengyo-Ando K, Kagawa-Nagamura Y, Kobayashi C, et al. Genetically encoded green fluorescent Ca^2+^ indicators with improved detectability for neuronal Ca^2+^ signals. PLoS One. 2012;7(12):e51286.

40. Lütcke H, Murayama M, Hahn T, Margolis DJ, Astori S, Zum Alten Borgloh SM, et al. Optical recording of neuronal activity with a genetically-encoded calcium indicator in anesthetized and freely moving mice. Front Neural Circuits. 2010;4:9.

41. Serrat R, Covelo A, Kouskoff V, Delcasso S, Ruiz-Calvo A, Chenouard N, et al. Astroglial ER-mitochondria calcium transfer mediates endocannabinoid-dependent synaptic integration. Cell Rep. 2021;37(12):110133.

42. Schindelin J, Arganda-Carreras I, Frise E, Kaynig V, Longair M, Pietzsch T, et al. Fiji: an open-source platform for biological-image analysis. Nat Methods. 2012;9(7):676–82.

43. Moreno-García Á, Bernal-Chico A, Colomer T, Rodríguez-Antigüedad A, Matute C, Mato S. Gene expression analysis of astrocyte and microglia endocannabinoid signaling during autoimmune demyelination. Biomolecules. 2020;10(9):1228.

44. Hou B, Zhang Y, Liang P, He Y, Peng B, Liu W, et al. Inhibition of the NLRP3-inflammasome prevents cognitive deficits in experimental autoimmune encephalomyelitis mice via the alteration of astrocyte phenotype. Cell Death Dis. 2020;11(5):377.

45. Liddelow SA, Guttenplan KA, Clarke LE, Bennett FC, Bohlen CJ, Schirmer L, et al. Neurotoxic reactive astrocytes are induced by activated microglia. Nature. 2017;541(7638):481–7.

46. Moreno-García A, Serrat R, Julio-Kalajzic F, Bernal-Chico A, Baraibar AM, Matute C, et al. In Vivo Assessment of cortical astrocyte network dysfunction during autoimmune demyelination: correlation with disease severity. J Neurochem. 2025;169(2):e16305.

47. Deshmukh VA, Tardif V, Lyssiotis CA, Green CC, Kerman B, Kim HJ, et al. A regenerative approach to the treatment of multiple sclerosis. Nature. 2013;502(7471):327–32.

48. Najm FJ, Madhavan M, Zaremba A, Shick E, Karl RT, Factor DC, et al. Drug-based modulation of endogenous stem cells promotes functional remyelination in vivo. Nature. 2015;522(7555):216–20.

49. Mei F, Lehmann-Horn K, Shen YA, Rankin KA, Stebbins KJ, Lorrain DS, et al. Accelerated remyelination during inflammatory demyelination prevents axonal loss and improves functional recovery. Elife. 2016;5.

50. Ransohoff RM. A polarizing question: do M1 and M2 microglia exist? Nat Neurosci. 2016;19(8):987–91.

51. Watanabe M, Toyama Y, Nishiyama A. Differentiation of proliferated NG2-positive glial progenitor cells in a remyelinating lesion. J Neurosci Res. 2002;69(6):826–36.

52. Moll NM, Hong E, Fauveau M, Naruse M, Kerninon C, Tepavcevic V, et al. SOX17 is expressed in regenerating oligodendrocytes in experimental models of demyelination and in multiple sclerosis. Glia. 2013;61(10):1659–72.

53. Bergner CG, van der Meer F, Franz J, Vakrakou A, Würfel T, Nessler S, et al. BCAS1-positive oligodendrocytes enable efficient cortical remyelination in multiple sclerosis. Brain. 2024;148(3):908–920.

54. Fard MK, van der Meer F, Sánchez P, Cantuti-Castelvetri L, Mandad S, Jäkel S, et al. BCAS1 expression defines a population of early myelinating oligodendrocytes in multiple sclerosis lesions. Sci Transl Med. 2017;9(419).

55. Moldrich G, Wenger T. Localization of the CB_1_ cannabinoid receptor in the rat brain. An immunohistochemical study. Peptides. 2000;21(11):1735–42.

56. Dudek KA, Paton SEJ, Binder LB, Collignon A, Dion-Albert L, Cadoret A, et al. Astrocytic cannabinoid receptor 1 promotes resilience by dampening stress-induced blood-brain barrier alterations. Nat Neurosci. 2025.

57. Yosef N, Xi Y, McCarty JH. Isolation and transcriptional characterization of mouse perivascular astrocytes. PLoS One. 2020;15(10):e0240035.

58. Rodriguez JJ, Mackie K, Pickel VM. Ultrastructural localization of the CB_1_ cannabinoid receptor in mu-opioid receptor patches of the rat Caudate putamen nucleus. J Neurosci. 2001;21(3):823–33.

59. Argaw AT, Asp L, Zhang J, Navrazhina K, Pham T, Mariani JN, et al. Astrocyte-derived VEGF-A drives blood-brain barrier disruption in CNS inflammatory disease. J Clin Invest. 2012;122(7):2454–68.

60. Chapouly C, Tadesse Argaw A, Horng S, Castro K, Zhang J, Asp L, et al. Astrocytic TYMP and VEGFA drive blood-brain barrier opening in inflammatory central nervous system lesions. Brain. 2015;138(Pt 6):1548–67.

61. Horng S, Therattil A, Moyon S, Gordon A, Kim K, Argaw AT, et al. Astrocytic tight junctions control inflammatory CNS lesion pathogenesis. J Clin Invest. 2017;127(8):3136–51.

62. Gimenez MA, Sim JE, Russell JH. TNFR1-dependent VCAM-1 expression by astrocytes exposes the CNS to destructive inflammation. J Neuroimmunol. 2004;151(1-2):116–25.

63. Williams JL, Manivasagam S, Smith BC, Sim J, Vollmer LL, Daniels BP, et al. Astrocyte-T cell crosstalk regulates region-specific neuroinflammation. Glia. 2020;68(7):1361–74.

64. Bullard DC, Hu X, Schoeb TR, Collins RG, Beaudet AL, Barnum SR. Intercellular adhesion molecule-1 expression is required on multiple cell types for the development of experimental autoimmune encephalomyelitis. J Immunol. 2007;178(2):851–7.

65. Héry C, Sébire G, Peudenier S, Tardieu M. Adhesion to human neurons and astrocytes of monocytes: the role of interaction of CR3 and ICAM-1 and modulation by cytokines. J Neuroimmunol. 1995;57(1-2):101–9.

66. Fernández-Moncada I, Lavanco G, Fundazuri UB, Bollmohr N, Mountadem S, Dalla Tor T, et al. A lactate-dependent shift of glycolysis mediates synaptic and cognitive processes in male mice. Nat Commun. 2024;15(1):6842.

67. Jimenez-Blasco D, Busquets-Garcia A, Hebert-Chatelain E, Serrat R, Vicente-Gutierrez C, Ioannidou C, et al. Glucose metabolism links astroglial mitochondria to cannabinoid effects. Nature. 2020;583(7817):603–8.

68. Argaw AT, Zhang Y, Snyder BJ, Zhao ML, Kopp N, Lee SC, et al. IL-1beta regulates blood-brain barrier permeability via reactivation of the hypoxia-angiogenesis program. J Immunol. 2006;177(8):5574–84.

69. Alvarez JI, Saint-Laurent O, Godschalk A, Terouz S, Briels C, Larouche S, et al. Focal disturbances in the blood-brain barrier are associated with formation of neuroinflammatory lesions. Neurobiol Dis. 2015;74:14–24.

70. Hind WH, Tufarelli C, Neophytou M, Anderson SI, England TJ, O’Sullivan SE. Endocannabinoids modulate human blood-brain barrier permeability in vitro. Br J Pharmacol. 2015;172(12):3015–27.

71. Panikashvili D, Shein NA, Mechoulam R, Trembovler V, Kohen R, Alexandrovich A, et al. The endocannabinoid 2-AG protects the blood-brain barrier after closed head injury and inhibits mRNA expression of proinflammatory cytokines. Neurobiol Dis. 2006;22(2):257–64.

72. Mestre L, Iñigo PM, Mecha M, Correa FG, Hernangómez-Herrero M, Loría F, et al. Anandamide inhibits Theiler’s virus induced VCAM-1 in brain endothelial cells and reduces leukocyte transmigration in a model of blood brain barrier by activation of CB_1_ receptors. J Neuroinflammation. 2011;8:102.

73. Liedtke W, Edelmann W, Chiu FC, Kucherlapati R, Raine CS. Experimental autoimmune encephalomyelitis in mice lacking glial fibrillary acidic protein is characterized by a more severe clinical course and an infiltrative central nervous system lesion. Am J Pathol. 1998;152(1):251–9.

74. Toft-Hansen H, Füchtbauer L, Owens T. Inhibition of reactive astrocytosis in established experimental autoimmune encephalomyelitis favors infiltration by myeloid cells over T cells and enhances severity of disease. Glia. 2011;59(1):166–76.

75. Voskuhl RR, Peterson RS, Song B, Ao Y, Morales LB, Tiwari-Woodruff S, et al. Reactive astrocytes form scar-like perivascular barriers to leukocytes during adaptive immune inflammation of the CNS. J Neurosci. 2009;29(37):11511–22.

76. Feliú A, Bonilla Del Río I, Carrillo-Salinas FJ, Hernández-Torres G, Mestre L, Puente N, et al. 2-Arachidonoylglycerol reduces proteoglycans and enhances remyelination in a progressive model of demyelination. J Neurosci. 2017;37(35):8385–98.

77. Guadalupi L, Mandolesi G, Vanni V, Balletta S, Caioli S, Pavlovic A, et al. Pharmacological blockade of 2-AG degradation ameliorates clinical, neuroinflammatory and synaptic alterations in experimental autoimmune encephalomyelitis. Neuropharmacology. 2024;252:109940.

78. Wang F, Han J, Higashimori H, Wang J, Liu J, Tong L, et al. Long-term depression induced by endogenous cannabinoids produces neuroprotection via astroglial CB. J Cereb Blood Flow Metab. 2019;39(6):1122–37.

