## Supplementary figure 1 for "Astrocyte CB_1_ receptors drive blood-brain barrier disruption in CNS inflammatory disease"

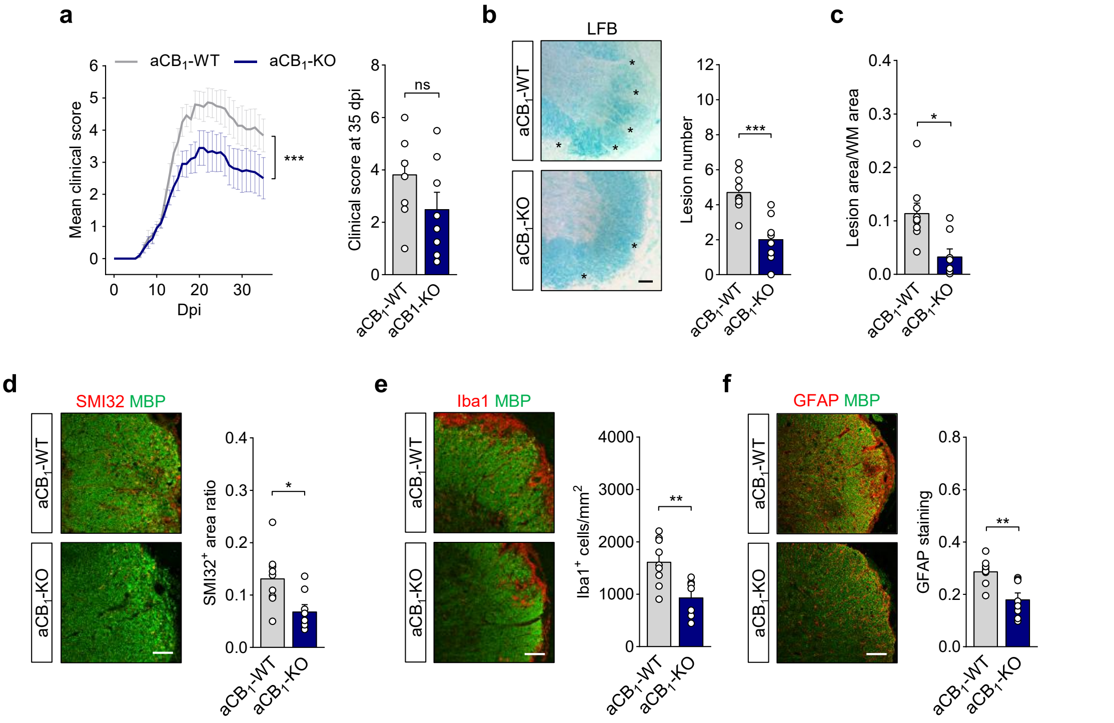


**Supplementary figure 1.** Reduced clinical severity and spinal cord pathology of chronic EAE in astrocyte-specific $CB_1$R null mice. (**a**) Clinical scores of aCB$_1$-KO and aCB$_1$-WT mice during chronic EAE. Comparison of motor scores from symptom onset to 35 dpi revealed significantly attenuated neurological disability in aCB$_1$-KO. Data are representative of 2 independent EAE experiments pooled together ($n$ = 8-12 mice). (**b**, **c**) Representative images of luxol fast blue myelin staining (LFB) and quantification of (**b**) demyelinating lesions (*) and (**c**) proportion of demyelinated area in spinal cord sections from aCB$_1$-KO and aCB$_1$-WT at 35 dpi ($n$ = 8-9 mice). Scale bar = 50 µm. (**d**-**f**) Immunohistochemical analysis of spinal cord inflammatory lesions at 35 dpi shows (**d**) reduced SMI32 positive profiles, (**e**) lower numbers of microglia/macrophages and (**f**) attenuated astrocyte reactivity in aCB$_1$-KO mice ($n$ = 7-9 mice). Scale bars = 100 µm. *$p < 0.05$, **$p < 0.01$ and ***$p < 0.001$, unpaired $t$-test or Mann-Whitney test.
