## Supplementary figure 2 for "Astrocyte CB_1_ receptors drive blood-brain barrier disruption in CNS inflammatory disease"

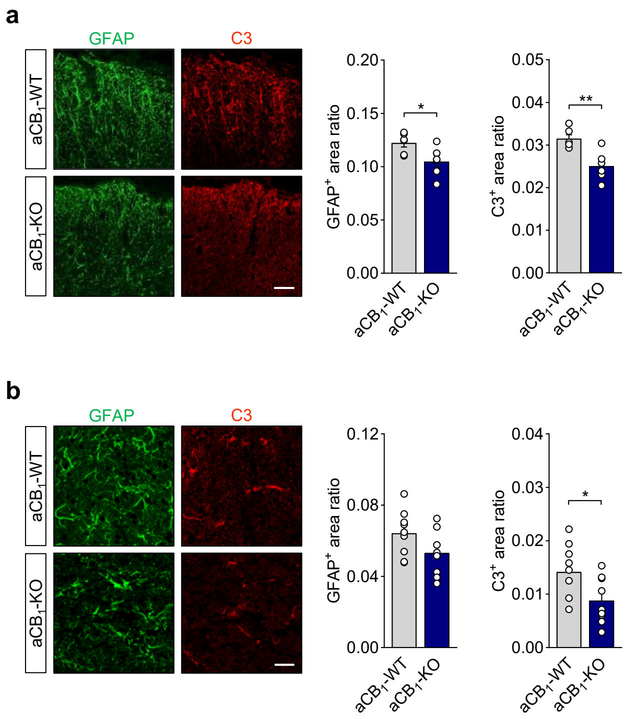


**Supplementary figure 2.** Restricted inflammatory neuropathology in aCB1-KO mice at acute EAE disease. Representative images and quantification of GFAP and C3 immunostaining in (**a**) spinal cord demyelinating lesions ($n$ = 6 mice) and (**b**) somatosensory cortex layers V-VI ($n$ = 8-9 mice) from in aCB1-KO and aCB1-WT mice at acute EAE disease normalized to tissue area. Scale bar = 100 µm (**a**) and 25 µm (**b**). *$p$ < 0.05 and **$p$ < 0.01, unpaired $t$-test or Mann-Whitney test.
