## Supplementary figure 3 for "Astrocyte CB_1_ receptors drive blood-brain barrier disruption in CNS inflammatory disease"

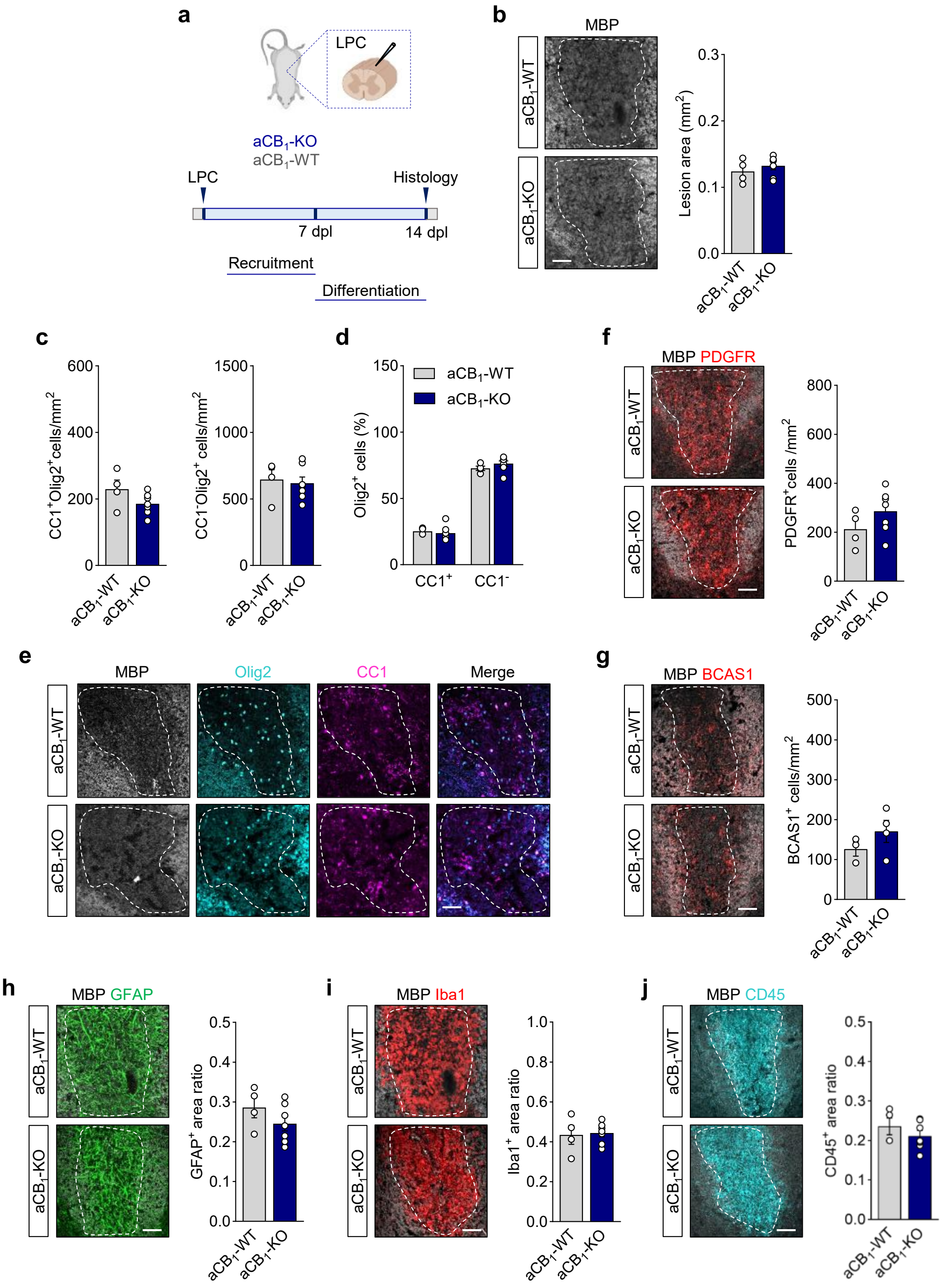


**Supplementary figure 3.** Astrocyte-encoded $CB_1R$ do not modulate oligodendrocyte populations in toxin induced remyelinating lesions. (**a**) Analysis of $aCB_1$-KO and $aCB_1$-WT mice in the LPC model of toxic demyelination. LPC lesions were analyzed at 14 dpl corresponding to oligodendrocyte differentiation and onset of remyelination. Data are representative of 2 independent LPC experiments pooled together. (**b**) Immunohistochemistry for MBP depicts demyelinated lesions (dashed lines) in the *dorsal funiculus* of LPC-injected mice. Scale bars = 50 μm. Quantitative analysis of lesion area shows no differences between genotypes (*n* = 4-7 mice). (**c**-**d**) Quantification of $CC1^+$/$Olig2^+$ oligodendrocytes and $CC1^-$/$Olig2^+$ immature oligodendroglia in LPC lesions indicates equal cell populations in $aCB_1$-KO and $aCB_1$-WT animals (*n* = 4-7 mice). Figure **e** shows representative micrographs of LPC lesions triple labelled for MBP, Olig2 and CC1. Scale bar = 50 μm. (**f**-**g**) Confocal images of LPC lesions double stained for MBP and (**f**) PDGFR or (**g**) BCAS1. Analysis of $PDGFR^+$ OPCs and $BCAS1^+$ myelinating oligodendrocytes in lesions from $aCB_1$-KO and $aCB_1$-WT mice shows no variations between genotypes (*n* = 3-7 mice). Scale bars = 50 μm. (**h**-**j**) Representative microphotographs and quantitative analysis of (**h**) astrocyte reactivity, (**i**) microglia/macrophage numbers and (**j**) in LPC lesions from $aCB_1$-KO and $aCB_1$-WT mice double labelled for MBP and GFAP, Iba1 or CD45. Scale bars = 100 μm.
