## Supplementary figure 4 for "Astrocyte CB_1_ receptors drive blood-brain barrier disruption in CNS inflammatory disease"

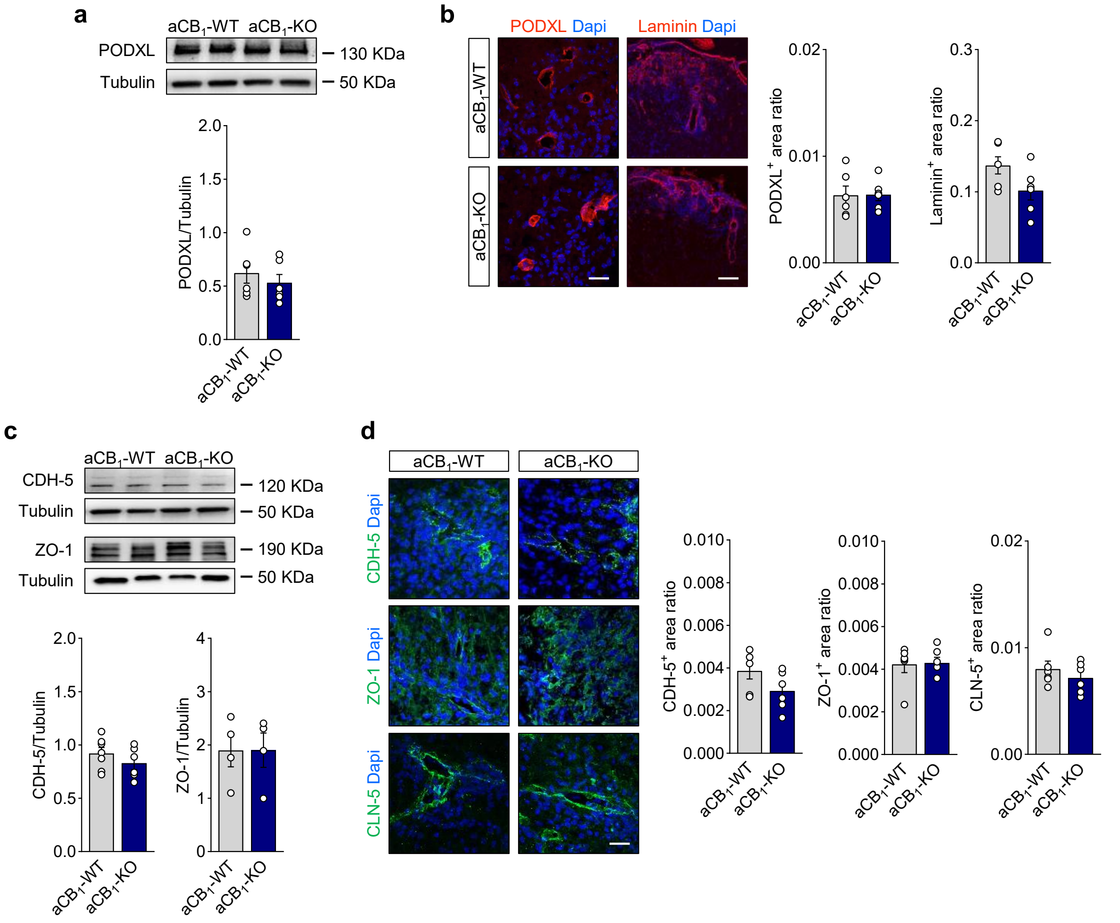


**Supplementary figure 4**. Analysis of the vascular endothelium in EAE lesions from mice lacking astrocyte-encoded $CB_1R$. (**a**, **b**) Immunoblotting and/or morphometry analysis of the endothelial cell markers PODXL and laminin in spinal cord tissue from $aCB_1$-KO mice and control $aCB_1$-WT mice at acute EAE disease (*n* = 6 mice). Scale bars = 50 μm (PODXL) and 100 μm (laminin). (**c**, **d**) Expression of endothelial TJ proteins CDH-5, ZO-1 and CLN-5 in spinal cord tissue from $aCB_1$-KO and $aCB_1$-WT mice was determined by immunoblotting (**c**) and/or immunofluorescence (**d**) (*n* = 6 mice). Scale bars = 25 μm.
