## Supplementary figure 5 for "Astrocyte CB_1_ receptors drive blood-brain barrier disruption in CNS inflammatory disease"

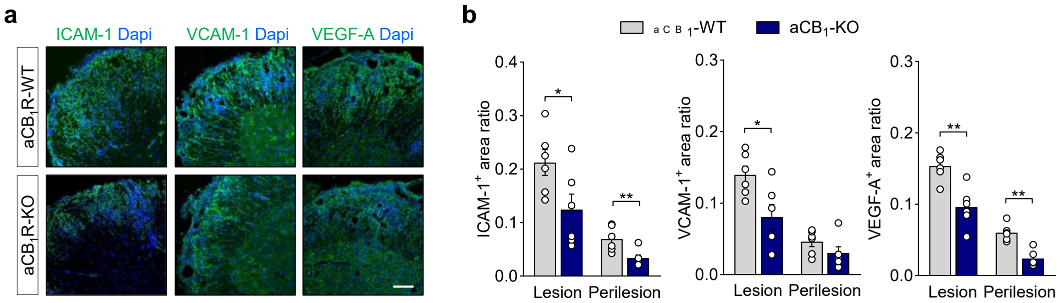


**Supplementary figure 5**. Reduced expression of vascular effector molecules in EAE lesions from astrocyte-specific $CB_1R$ null mice. Representative confocal micrographs (**a**) and morphometry (**b**) show lower expression levels of the adhesion molecules ICAM-1 and VCAM-1 and VEGF-A within lesion (L) and perilesion (PL) areas from aCB1-KO mice (*n* = 6 mice). Scale bar = 100 μm. **p* < 0.05 and ***p* < 0.01, unpaired *t*-test or Mann-Whitney test.
