## Supplementary Table 1 for "Astrocyte CB_1_ receptors drive blood-brain barrier disruption in CNS inflammatory disease"

**Supplementary Table 1. Primers for RT-qPCR analysis**

| Gene | Forward primer | Reverse primer |
|---|---|---|
| *B2m* | ACTGACCGGCCTGTATGCTA | ATGTTCGGCTTCCCATTCTCC |
| *Bdnf* | TCCAAAGGCCAACTGAAGCA | CTGCAGCCTTCCTTGGTGTA |
| *C1ra* | CCTTCCACCCCATAGAACAG | CCTTCAACTTACAGTGCCTGA |
| *C3* | AGGGAGTGTTTGTGCTGAAC | GCCAATGTCTGCCTTCTCTAC |
| *Ccl2* | GTAGTTTTTGTCACCAAGCTC | CAGAAGTGCTTGAGGTGGTT |
| *Ccl5* | GCTCCAATCTTGCAGTCGT | CCTCTATCCTAGCTCATCTCCA |
| *Cldn1* | ACAGCATGGTATGGAAACAGA | AGGAGCAGGAAAGTAGGACA |
| *Cldn4* | CACTCAGCCTACACGTTACTC | CACTCAGCACACCATGACTT |
| *Cntf* | GTGAAGACAGAAGCAAACCAG | AGATAGAGCGGCTACAGA GG |
| *Cxcl10* | ATTTTCTGCCTCATCCTGCT | TGATTTCAAGCTTCCCTATGGC |
| *Fkbp5* | GACACCAAAGAAAAGCTGACG | ACTATCTTCCTGTACTGA ATC |
| *Gapdh* | AGACGGCCGCATCTTCTT | TTCACACCGACCTTCACCAT |
| *Gbp2* | GCAGAATTCACCTCATACATCTTG | GATGGCACCAACATAGGTCTG |
| *Ggta1* | TCCTGTTGATGCTGATTGTCTC | GTCCTTCTGCCATCTGTT CTC |
| *H2-T23* | ACAGTCCCGACCCAGAGTAG | CCACGTAGCCGACAATGATGA |
| *Hprt* | CAGTACAGCCCCAAAATGGTTA | AGTCTGGCCTGTATCCAACA |
| *Icam1* | CTGTGCTTTGAGAACTGTGG | GGTCCTTGCCTACTTGCTG |
| *Igf1* | ATGCTCTTCAGTTCGTGTGT | AGTACATCTCCAGTCTCC TCA G |
| *Iigp1* | CTCATGTGAAGAGCCTGTAGC | CTGACCCATGACTTCAAGCA |
| *Mbp* | CCCTCACAGCGATCCAAGTA | CTCTGTGCCTTGGGAGGAA |
| *Mog* | CACTTGTGCCTACGATCCTC | AGTCCGATGGAGATTCTCTACT |
| *Ntf3* | AGTCCACCTTTCTCTTCATGTC | ACATCACCTTGTTCACCTGTA |
| *Olig2* | AGGGATGATCTAAGCTCTCGAA | ATTACAGACCGAGCCAAC AC |
| *Pdgfa* | ATTAACCATGTGCCCGAGA | GTATCTCGTAAATGACCGTCC |
| *Pdgfra* | TCACAGCCACCTTCATTACAG | GTTGCCTTACGACTCCAGATG |
| *Ppia* | AGGGTTCCTCCTTTCACAGAA | TGCCGCCAGTGCCATTA |
| *Psmb8* | GCTGCTTTCCAACATGATGC | CCGAGTCCCATTGTCATC TAC |
| *Serping* | CTACCAAGATGGCTAAGACCAA | GATGCTCTCCAAGTTGCT CT |
| *Tgfb1* | GCTGCGCTTGCAGAGATTAA | GTAACGCCAGGAATTGTTGCTA |
| *Vcam1* | GCAAAGGACACTGGAAAAGAG | TGTGCAGTTGACAGTGACA |
| *Vegfa* | AGAAAGACAGAACAAAGCCAGA | TCGTTTAACTCAAGCTGCCT |
