## Supplementary Table 2 for "Astrocyte CB_1_ receptors drive blood-brain barrier disruption in CNS inflammatory disease"

**Supplementary Table 2. Primary antibodies for immunohistochemistry**

| Antibody | Host | Dilution | Reference |
|---|---|---|---|
| B220 | Rat | 1:200 | BD Biosciences, 557390 |
| BCAS1 | Rabbit | 1:500 | Synaptic Systems, 445003 |
| C3 | Rabbit | 1:1000 | Dako, A0063 |
| CDH-5 | Goat | 1:100 | R&D Systems, AF1002 |
| CC1 | Mouse | 1:200 | Calbiochem, OP80 |
| CCL2 | Mouse | 1:300 | Santa Cruz, sc-32771 |
| CD3 | Rat | 1:50 | Bio-Rad, MCA1477 |
| CD45 | Rat | 1:100 | BD Pharmigen, 550539 |
| CD68 | Rat | 1:100 | Bio-Rad, MCA1957GA |
| CLN-1 | Mouse | 1:100 | ThermoFisher, 37-4900 |
| CLN-4 | Mouse | 1:100 | ThermoFisher, 32-9400 |
| CLN-5 | Mouse | 1:100 | Inivtrogen, 35-2500 |
| Fibrinogen | Sheep | 1:200 | US Biological, F4199 |
| GFAP | Rabbit | 1:2000 | Dako, Z0334 |
| GFAP | Chicken | 1:1000 | Abcam, Ab4674 |
| Iba1 | Rabbit | 1:500 | Wako, 019-19741 |
| ICAM-1 | Goat | 1:500 | R&D Systems, AF796 |
| Laminin | Rabbit | 1:500 | Merck, L9393 |
| Ly6G | Rat | 1:100 | BioLegend, 127601 |
| MBP | Chicken | 1:200 | Merck, ab9348 |
| MBP | Mouse | 1:1000 | Biolegend, 808402 |
| NG2 | Rabbit | 1:200 | Merck, AB5320 |
| Olig2 | Mouse | 1:500 | Merck, MABN50 |
| PDGFRα | Goat | 1:250 | R&D Systems, AF1062 |
| PODXL | Goat | 1:500 | R&D Systems, AF1556 |
| SMI32 | Mouse | 1:200 | BioLegend, 801702 |
| VCAM-1 | Rabbit | 1:500 | Abcam, ab134047 |
| VEGF-A | Rabbit | 1:500 | Santa cruz, sc-507 |
| ZO-1 | Rabbit | 1:100 | Invitrogen, 402200 |
